# Dynamic de novo adipose tissue development during metamorphosis in *Drosophila melanogaster*

**DOI:** 10.1101/2022.03.29.486213

**Authors:** Taiichi Tsuyama, Hanae Komai, Yusaku Hayashi, Kohei Shimono, Tadashi Uemura

**Affiliations:** Graduate School of Biostudies, Kyoto University, Kyoto 606-8501, Japan; Research Center for Dynamic Living Systems, Kyoto University, Kyoto 606-8501, Japan; AMED-CREST, AMED, 1-7-1 Otemachi, Chiyoda-ku, Tokyo 100-0004, Japan

**Keywords:** Insect development, adult fat body, adipose tissue, directional migration, cell proliferation, in vivo imaging

## Abstract

Adipose tissue is a central organ for controlling systemic metabolism both in invertebrates and vertebrates. Here, we have investigated the cellular mechanisms of the adult-type fat body (AFB) development in *Drosophila*. We have established genetic tools that allow visualization and genetic manipulations of cells in the AFB lineage from early in metamorphosis. We identified precursor cells that give rise to the AFB and delineated dynamic cellular mechanisms underlying AFB formation. These precursor cells displayed polarized cell shapes and oriented motility, with emigration from the thorax and subsequent dispersal to the abdomen and head. After the migration period, these cells adhered to each other, assembling into the AFB with a sheet-like architecture. Continuous cell proliferation occurred during and after the large-scale migration to achieve appropriate fat tissue mass. Homotypic cell fusion after the sheet formation contributed to the establishment of multinucleated cells in the AFB. We also examined candidate gene functions, and our results argue that Rac1, ecdysone signaling, and the transcription factor Serpent support adult fat body organogenesis.

**Brief Summary Statement:** *Drosophila* adult fat body precursor cells form adult adipose tissue during metamorphosis by directional migration, continuous cell proliferation, and homotypic cell fusion.

## Introduction

The adipose tissue is a central organ for the control of systemic metabolism across various animal orders, including invertebrates and vertebrates (Gesta et al., 2007). It plays essential roles in storing and mobilizing energy substrates and functions as a pivotal signaling center for inter-organ communications that regulate energetic metabolism, systemic growth, and immune responses at the organismal level (Droujinine and Perrimon, 2016; Rosen and Spiegelman, 2014). The adipose tissue predominantly, but not exclusively, comprises adipocytes, which accumulate large lipid droplets as energy storage. Adipocytes and other cell types are enclosed by layers of a basal lamina, thus assembled into a discrete organ. While molecular mechanisms controlling homeostasis in mature adipose tissue have been extensively studied, much less is known about developmental mechanisms underlying adipose tissue organogenesis (Sebo and Rodeheffer, 2019).

The adipose tissue correlate in insects is the fat body (Buys, 1924), which accumulates not only lipid droplets but also glycogen and protein granules and serves a wide array of physiological activities, including systemic control of metabolism and growth, innate immunity, and adult courtship behaviors (Arrese and Soulages, 2010; Boulan et al., 2015; Franz, et al., 2018; Li et al., 2019). *Drosophila* (*Sophophora*) *melanogaster* develops two distinct fat body types; the larval fat body (LFB) and adult fat body (AFB), both of which are solely composed of fat cells. The LFB develops during embryogenesis and persists until 2-3 days after adult eclosion, serving as a major energy reservoir in feeding larvae and as a nutriment reserve during metamorphosis and in newly eclosed adult flies (Aguila et al., 2007; Aguila et al., 2013; Butterworth et al., 1988). The AFB is found in the newly eclosed imago and functions during adult life (Johnson and Butterworth, 1985). The LFB and AFB are distributed throughout the body and reside in the body cavity between the gut and the body wall (Fig. S1A,B; Miller 1950; Rizki et al., 1980). Fat cells are ensheathed by connective tissues that form a characteristic one-cell thick sheet-like structure (Dai et al., 2017). Both types of *Drosophila* fat body have served as model systems to study the significance of flexible lipid mobilization and impacts of deregulated energetic metabolism in the face of various types of physiological or pathological perturbations (Alfa and Kim, 2016; Baumbach et al., 2014; DiAngelo and Birnbaum, 2009; Storelli et al., 2019).

Our understanding of developmental processes underlying fat body organogenesis has been provided chiefly by studies of LFB development in *Drosophila melanogaster* (Hoshizaki, 2005). While it is widely accepted that both LFB and AFB are mesodermal in origin, they arise from distinct cell lineages (Lawrence and Johnston, 1986). The larval fat cells are specified by positional information in the embryo, leading to the formation of fat cell progenitors in a metameric manner (Azpiazu et al., 1996; Riechmann et al., 1998). Histochemical studies showed that these progenitors proliferate to form clusters and are integrated into a continuous tissue covering a broad region of the larval body (Miller et al., 2002). Recent studies also shed light on many aspects of mechanisms underlying the establishment of tissue architectures and intracellular structural organizations of the LFB cells (Dai et al., 2018; Diaconeasa et al., 2013; Ugrankar, 2019; Zang, Wan et al., 2015). In contrast to the LFB, the early developmental trajectories giving rise to the AFB in *Drosophila* have been mostly unexplored for decades, except for a few studies (Ferrus and Kankel, 1981; Hoshizaki et al., 1995; Lawrence and Johnston, 1986).

Here, we have developed and combined genetic tools that allow Gal4 expression in the AFB lineage during metamorphosis, enabling us to label precursor cells giving rise to the AFB and to manipulate gene functions in these cells. The precursor cells emerged from central segments, dispersed to the abdomen and head, and assembled into the mature AFB. At the cellular level, this transformation involved directional migration, continuous proliferation, and homotypic cell fusion. At the genetic level, some of the key events in AFB organogenesis appeared to be governed by the Rac1 small GTPase, ecdysone signaling, and the Serpent transcription factor.

## Results

### Specific Gal4 labeling allows visualization of the developing adult fat body during metamorphosis

The AFB is found in newly eclosed adult flies (Johnson and Butturworth, 1985) but not in mature third instar larvae. Thus, major developmental events related to the AFB must take place during metamorphosis. In this study, we mainly focused on the abdomen, in which the AFB predominantly resides (Fig. 1 A; Miller, 1950). We first tested whether known fat body Gal4 drivers were able to highlight the AFB in young adult flies (0-1 day after adult eclosion; see Materials and Methods); however, all of the Gal4 lines that we tested drove GFP expression in the LFB, thus obscuring AFB development due to close proximity between the LFB and AFB. While larval fat cells undergo remodeling, involving autophagy induction and tissue dissociation, with progressive cell death during metamorphosis (Butterworth et al., 1988; Jia et al., 2017; Liu et al., 2013; Rusten et al., 2004), a large population of larval fat cells is persistent even at adult eclosion. To overcome this limitation, we adopted two strategies. First, we took advantage of the Gal80 protein, which inhibits Gal4 (Ma and Ptashne, 1987; Pfeiffer et al., 2010), in conjunction with cis-elements of genes that are expressed in the later development of the LFB (Fig. S1D). Second, we tested Gal4 lines associated with genes controlling the early development of the LFB.

**Fig. 1.**
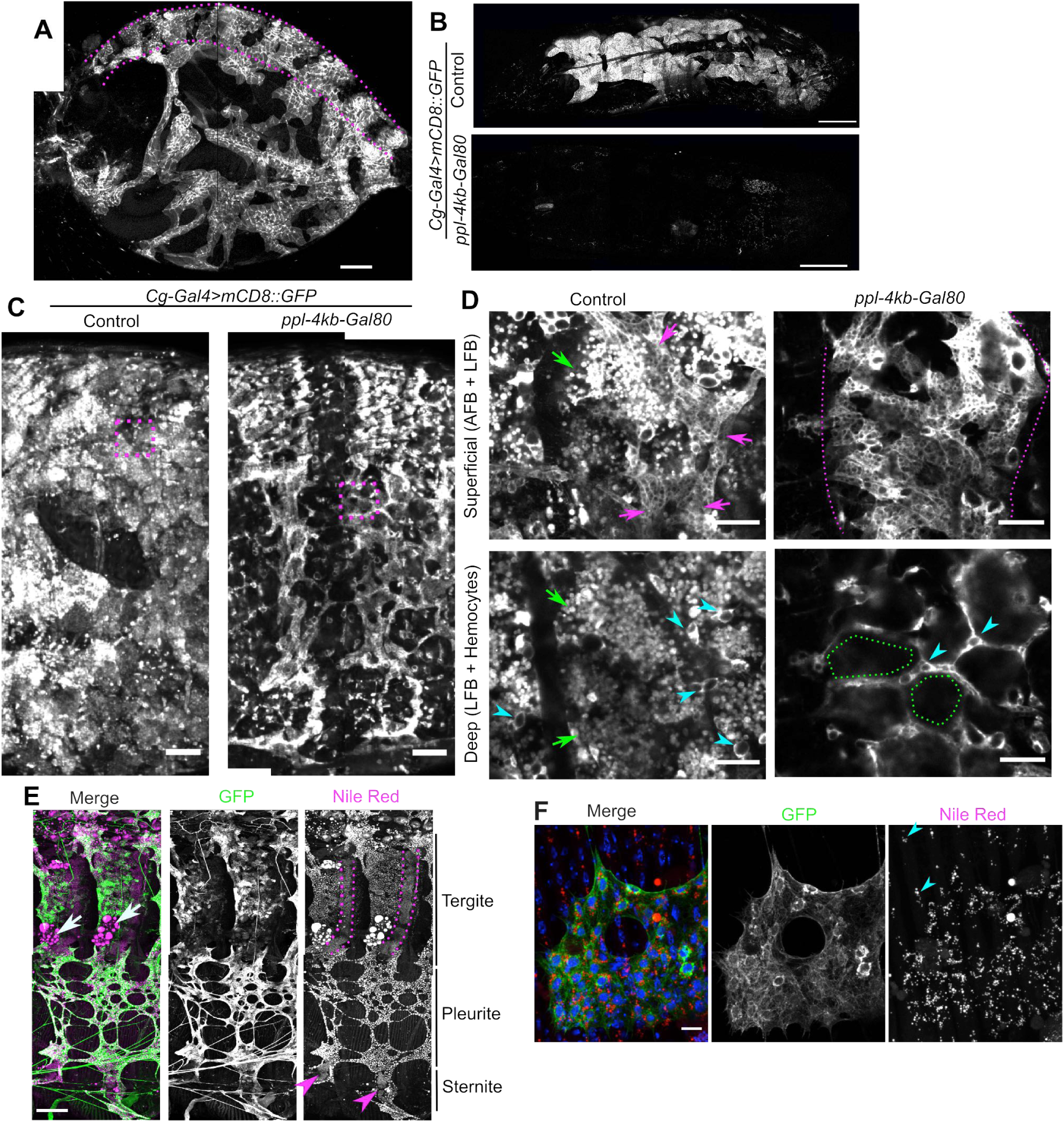
Gal4-based cell labeling highlights the adult fat body during metamorphosis. (A) AFB in the abdomen cut from a female fly (one-week-old). The AFB was visualized using *Cg-Gal4*. In all figures, cell membranes were visualized by membrane targeted fluorescent markers. The region between dashed lines indicates the tergite. (B-D) *ppl-4kb-Gal80* potently suppressed *Cg-Gal4* dependent GFP expression in the LFB from the mature 3rd instar stage to the late pharate adult stage. (B) Mature larvae expressing mCD8::GFP with the absence (top) or presence (bottom) of *ppl-4kb-Gal80*. (C,D) *ppl-4kb-Gal80* highlighted the abdominal AFB (C, right), which was concealed by strong GFP signals in larval fat cells in the absence of *ppl-4kb-Gal80* (C, left), in pharate adult flies shortly before eclosion. (D) High magnification optical slices at regions indicated by magenta rectangles in the low magnification z-stacked images (C). Superficial layers were 8 μm above deeper slices. The AFB was seen only at superficial layers (top; magenta arrows and the area between magenta dashed lines). Larval fat cells were seen both at superficial and deep layers, with membrane markers showing their foamy surface (green arrows; Diaconeasa et al., 2013). Hemocytes were predominantly seen only at the deep layers and collectively wrapped around larval fat cells (light blue arrowheads; Rizki, 1980), revealing the outlines of larval fat cells, in which GFP signals were diminished by *ppl-4kb-Gal80* (bottom right; green dashed lines). (E) GFP signals driven by *c833-Gal4* plus *ppl-4kb-Gal80* were detected in most Nile Red positive adult fat cells in the abdomen of a dissected pharate adult fly (70-75 h APF). White arrows in the merged channel indicate larval fat cells. Magenta dashed lines and arrowheads indicate dorsal oenocyte belts and ventral oenocyte clusters stained with Nile Red, respectively. (F) Numerous lipid droplets were detected in the AFB expressing GFP in the pleurite as early as 66 h APF, visualized with Nile Red for lipids (red) and Hoechst 33342 for nuclei (blue). Light blue arrowheads indicate lipid depositions in adult lateral muscle cells. Scale (um): 200 (A), 500 (B), 100 (C,E), 50 (D), 10 (F). In all images including subsequent figures, anterior is to the left. In all lateral view images, dorsal is up. Fly strains and exact genotypes are summarized in Tables 16 and 17, respectively.

We generated fly lines with Gal80 under the control of several cis-elements of fat-body-related genes (Fig. S1E,F, S2A-D, Table 1) and tested whether these transgenes could inhibit GFP expression in the LFB induced by *Cg-Gal4*, which is expressed in the fat body and hemocytes (Asha et al., 2003). Among these Gal80 strains, we focused on the *Gal80* gene induced by a cis-element of the *ppl* gene (hereafter referred to as *ppl-4kb-Gal80*; Buch et al., 2008; Zinke et al., 1999). *ppl-4kb-Gal80* potently decreased GFP signals in the LFB at the late third instar stage (Fig. 1B). The suppression of Gal4 activity in the LFB persisted until the late pharate adult stage, allowing the visualization of the AFB (Fig1. C,D). When a GFP marker was driven by *ppl-4kb-Gal80* and *c833-Gal4* (a known fat body Gal4 driver, Hrdlicka et al., 2002), tissues with a sheet-like architecture were seen in the abdomen from 60-65 h after puparium formation (APF) onward (flies were grown at 25°C unless otherwise noted). The tissues were efficiently stained with Nile Red, a neutral lipid marker, indicating that they indeed comprised the AFB (Fig. 1E). The accumulation of numerous lipid droplets in the AFB was detected as early as 66 h APF (Fig. 1F). At 60 h APF, GFP positive cells contained only tiny lipid droplets (data not shown), suggesting that the AFB starts to deposit lipids at 60-65 h APF. It is noteworthy that the specificity of this Gal4/Gal80 combination is rather limited. *c833-Gal4* expresses is expressed in some neurons, including peripheral sensory Class IV neurons (Grueber et al., 2002; Shimono, Fujimoto et al., 2009), and in surrounding tissues, including part of the adult epidermis, as revealed by the expression of nuclear-localized fluorescent protein markers (Fig. S2 E). In addition, *ppl-4kb-Gal80* suppressed Gal4-driven gene expression in hemocytes at the third instar and white prepupa stage (Fig. S3).

We also examined strains in Gal4 libraries driven by short cis-elements encoded in the fly genome (Jenett et al., 2012; Kvon et al., 2014). *seven-up* (*svp*) is expressed during the early development of the LFB (Hoshizaki et al., 1994). Two *svp* Gal4 strains (*svp[31F09]-Gal4* and *svp[31H02]-Gal4)* exhibited transient Gal4 expression early in the AFB lineage, as revealed by the G-TRACE system (Fig. S2F; Evans et al., 2009). To sustain the transient Gal4 expression induced by *svp[31F09]-Gal4*, we employed the *Ay-Gal4* system, a lineage tracing method (Fig. S2G; Ito et al., 1997). This *svp[31F09]-Gal4* plus *Ay-Gal4* combination drove UAS-dependent markers in the AFB more specifically than the *c833-Gal4* plus *ppl-4kb-Gal80* combination, although it induced Gal4-dependent marker expression in a small population of cells that moved with weak directionality, possibly hemocytes, and a few adult nephrocytes in the abdomen (Fig. S4D; Na et al., 2015).

While observing the AFB labeled with these Gal4-based tools, we noticed that various genotypes of flies expressing only fluorescent markers exhibited reduced or absent AFB signals with variable degrees of penetrance (Table 2; Fig. S4A-C). Those adult flies with diminished marker signals displayed reduced or missing Nile Red signals (N =11; Fig. S4D,E), suggesting that AFB development was disrupted in these flies. Unexpectedly, even *Canton-S*, a widely used wild-type strain, but not *Oregon-R*, showed disorganized or lost AFB tissues with approximately 30% penetrance (Fig. S4F,G, Table 3). To examine whether defects in AFB development are common in wild-type strains, we tested 39 strains in Drosophila Genetic Reference Panel, which were established as inbred lines from the Raleigh, USA population (Huang, Massouras et al., 2014). These strains rarely showed AFB defects, with 1.4% penetrance on average (N for each line was approximately 140; Table 4). Thus, the aberrant AFB development observed in wild-type genetic backgrounds might be due to genetic variations that might arise after establishment as laboratory strains or possibly cellular toxicity due to fluorescent protein expression (Ansari et al., 2016; Kintaka et al., 2016).

Taken together, we established genetic combinations that predominantly express Gal4 in the developing AFB, which starts to accumulate lipid droplets at 60-65 h APF.

### Adult fat body precursor cells migrate directionally over long-distance in the abdomen

The genetic tools described above enabled us to follow cells giving rise to the AFB as early as 15 h APF, revealing their directional migration and continuous proliferation. Since these cells were located beneath the integument in a quasi-two-dimensional manner (Fig. S5A,B, Movie 1), their developmental dynamics were readily observed using live imaging without dissection. We refer to these cells prior to the AFB formation (approximately 65 h APF) as AFB precursor (AFBp) cells hereafter. These cells extended and retracted many prominent filopodia-like and lamellipodia-like protrusions, which may support their directional and persistent motility (Fig. 2A, Movie 2).

**Fig. 2.**
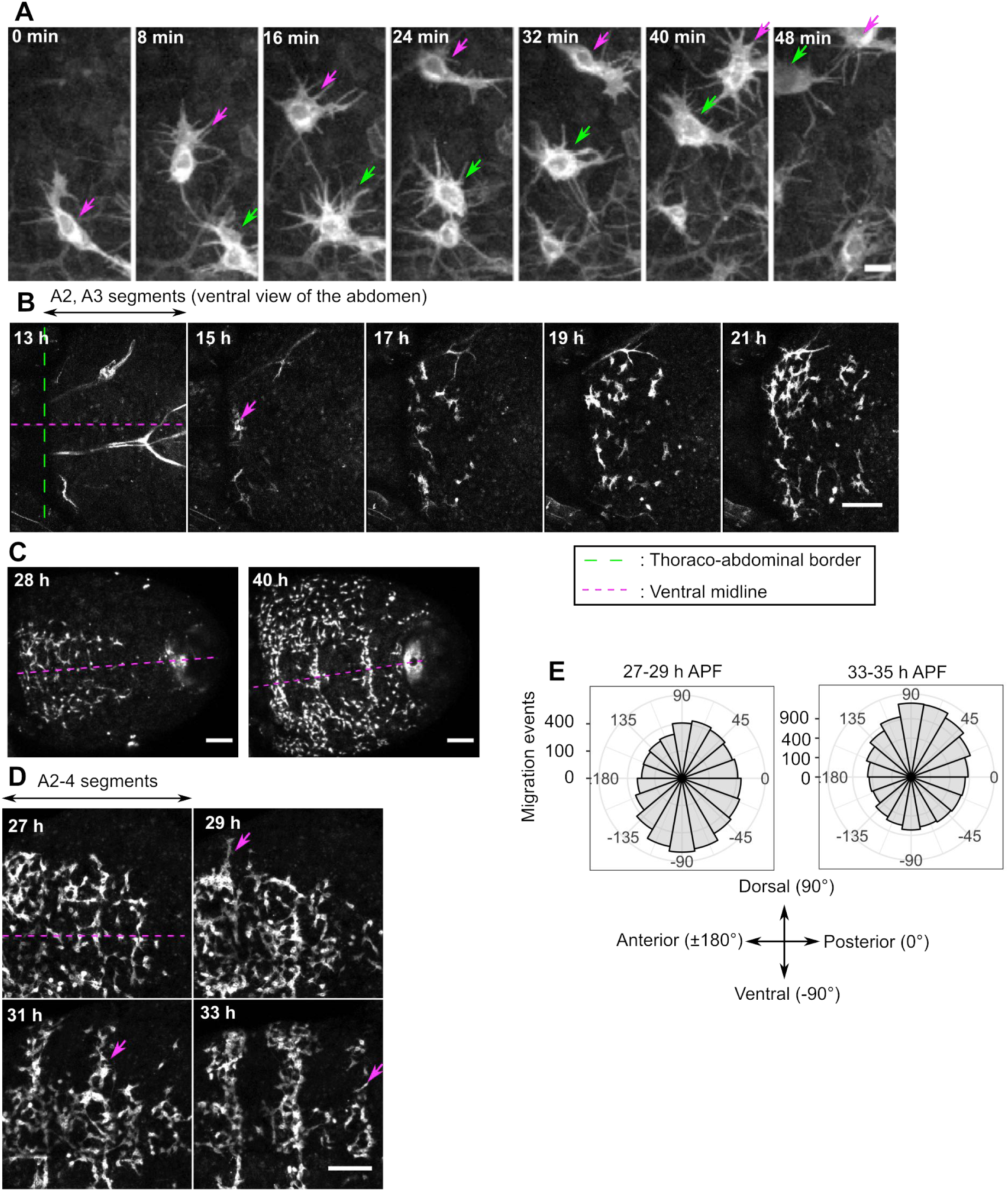
AFBp cells emerge from the ventroanterior end of the abdomen and migrate posteriorly. (A) Actively migrating AFBp cells showing cellular protrusions (34 h APF). Magenta and green arrows indicate the same cells over time. (B) Ventral view image sequences of the anterior part of the abdomen. Green and magenta dashed lines indicate the thoraco-abdominal border and the VML, respectively. AFBp cells were first seen at 15 h APF (magenta arrow) and then migrated posteriorly. Axons extending from the CNS were seen at 13 h and degenerated by 15 h APF. (C) AFBp cells dispersed over the ventral aspect of the abdomen by 40 h APF. (D) Ventral AFBp cells started to migrate dorsally around 30 h APF (magenta arrows). (E) Quantification of migration direction of AFBp cells labeled with nuclear-targeted GFP (5168 and 9384 steps greater than 1 μm/min from 4 female flies for 27-29 and 33-35 h, respectively). Rose diagrams show migration step numbers in each direction bin (bin size: 20°). The area size but not the height of each sector indicates step numbers. Scale (μm): 10 (A), 100 (B-D).

In the abdomen, AFBp cells were first seen at the ventroanterior end at approximately 15 h APF, from which site they migrated posteriorly and eventually reached the posterior end, covering almost the entire area of the ventral aspect of the abdomen by 40 h (Fig. 2B,C, Movie 3, 4). These data suggest that AFBp cells originate and emigrate from more anterior segments, the thorax or head. At approximately 30 h APF, a subset of ventral AFBp cells started to migrate dorsally, entering pleurites (Fig. 2D). The onset timing of the dorsal switching differed along the anteroposterior (AP) axis, with AFBp cells in anterior segments starting the dorsal migration earlier. To quantitatively analyze their migration patterns, we tracked AFBp cells during 27-29 and 33-35 h APF and measured angle and displacement values of nuclear movements at 2 min intervals (Fig. 2E, Movie 5). At 27-29 h APF, their migration steps were biased toward the ventroposterior direction. At 33-35 h APF, their migration direction reversed along the dorsoventral (DV) axis (Fig. 2E, Table 5). AFBp cells distant from the ventral midline (VML) more frequently moved dorsally than those in other regions at 33-35 h APF, but not at 27-29 h APF (Fig. S5C). Both at 27-29 and 33-35 h APF, AFBp cell migration along the AP axis was biased toward the posterior direction (Fig. 2E). We also measured displacement values of migration events subdivided by four directional categories (dorsal, ventral, anterior, and posterior) and found that anterior movements were slightly smaller than migrations toward other directions (Fig. S5 D, Table 6). These data suggest that the directional bias toward the posterior direction, rather than differences in the displacement length of movements toward each direction, is a significant driver of the posterior migration of AFBp cells.

The observed directional tendency toward the VML at earlier times (27-29 h) might be imposed by rearrangements of surrounding tissues. From 25 to 35 h APF, the length of the migration area along the DV axis progressively decreased (S5E,F, Movie 6), possibly suggesting that other tissues physically confine the migration area for AFBp cells.

As described above, dorsally migrating AFBp cells proceeded through a narrow path in each hemisegment and entered the pleurites. Leading AFBp cells passed the abdominal spiracle at 40 h (Fig. 3A, Movie 7) and reached the dorsal midline by 50 h APF (Fig. 3B, Movie 8). The directional bias of AFBp cell migration at the pleurites varied over time. A gradual reversal in their migration direction occurred around abdominal spiracles during 47-53 h APF (Fig. 3C,D, Movie 9, Table 7). The displacement length of dorsal migration appeared to decrease during the same period (Fig. S5G, Table 8). In contrast to AFBp cells, LFB cells and hemocytes in the same region displayed only small movements without directional bias (Movie 10).

**Fig. 3.**
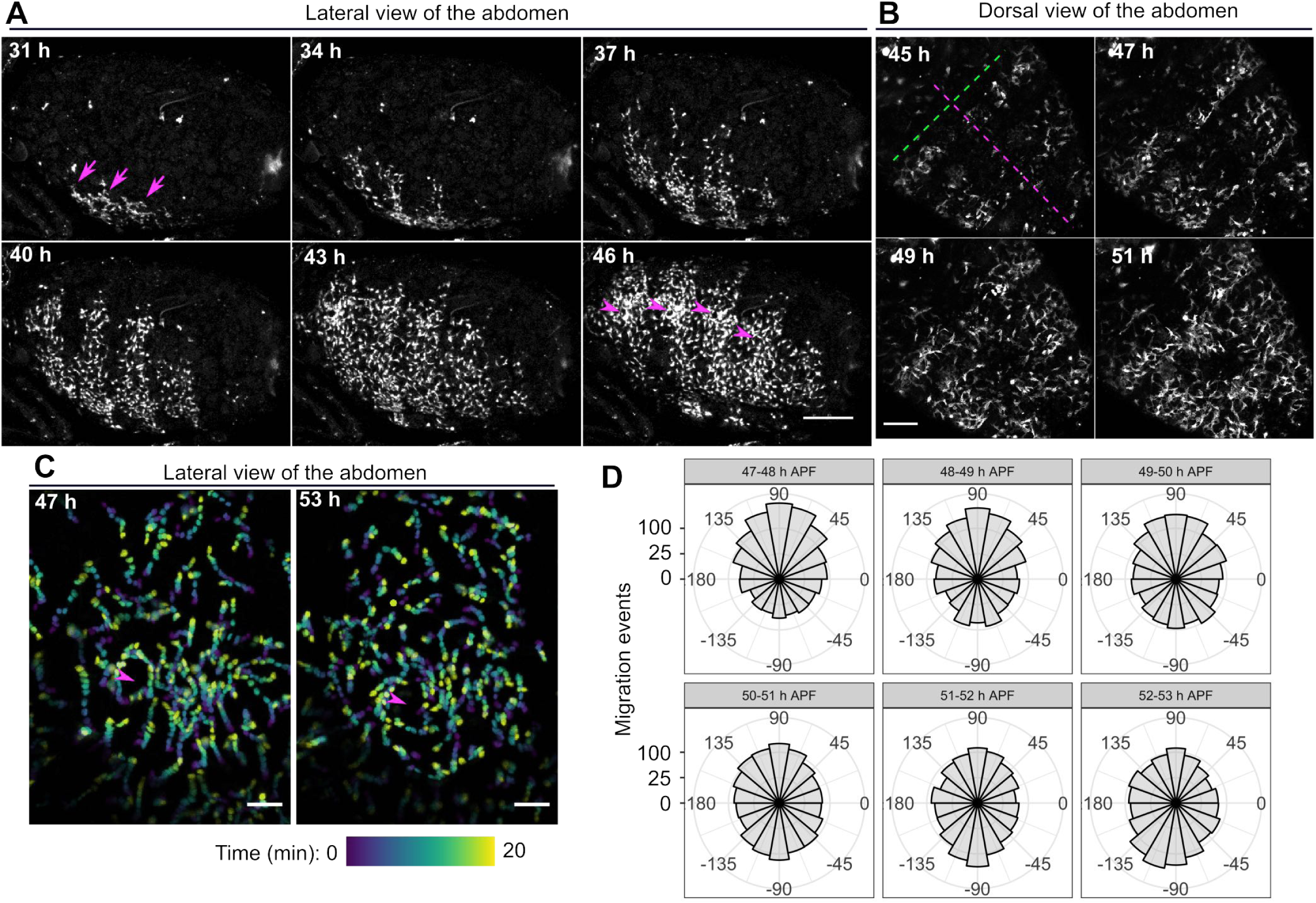
AFBp cells from the ventral side cover the pleurite and tergite by dorsal migration. (A) Lateral view image stills of abdominal AFBp cells entering the pleurite. Arrows and arrowheads indicate the positions of AFBp cells starting dorsal migration and the abdominal spiracles, respectively. (B) Dorsal view image stills of abdominal AFBp cells covering the tergite. Dashed lines indicate the dorsal midline (magenta) and the thoraco-abdominal border (green), respectively. (C) Representative temporal-color coded images revealed a directional switching in the pleurite. Arrowheads indicate the abdominal spiracle. AFBp cells expressing nuclear-targeted GFP were imaged over time. Elapsed time after the reference time (47 or 53 h APF) is color-coded, as indicated by the color code below. (D) Quantification of migration direction of AFBp cells around abdominal spiracles (7049 steps in total from 4 female flies). Scale (μm), 100 (A,B), 50 (C).

The characteristic sheet-like architecture of the AFB was achieved by 65 h APF across all abdominal regions (Fig. 4A,B, Movie 11). After the migration period, AFBp cells became flattened but continued to extend fine filopodia-like protrusions in various directions (Fig. 4C, Movie 12). These cells increased contact areas with neighboring cells as fine protrusions were retracted. Eventually, the characteristic monolayer was established (Fig. 4D).

**Fig. 4.**
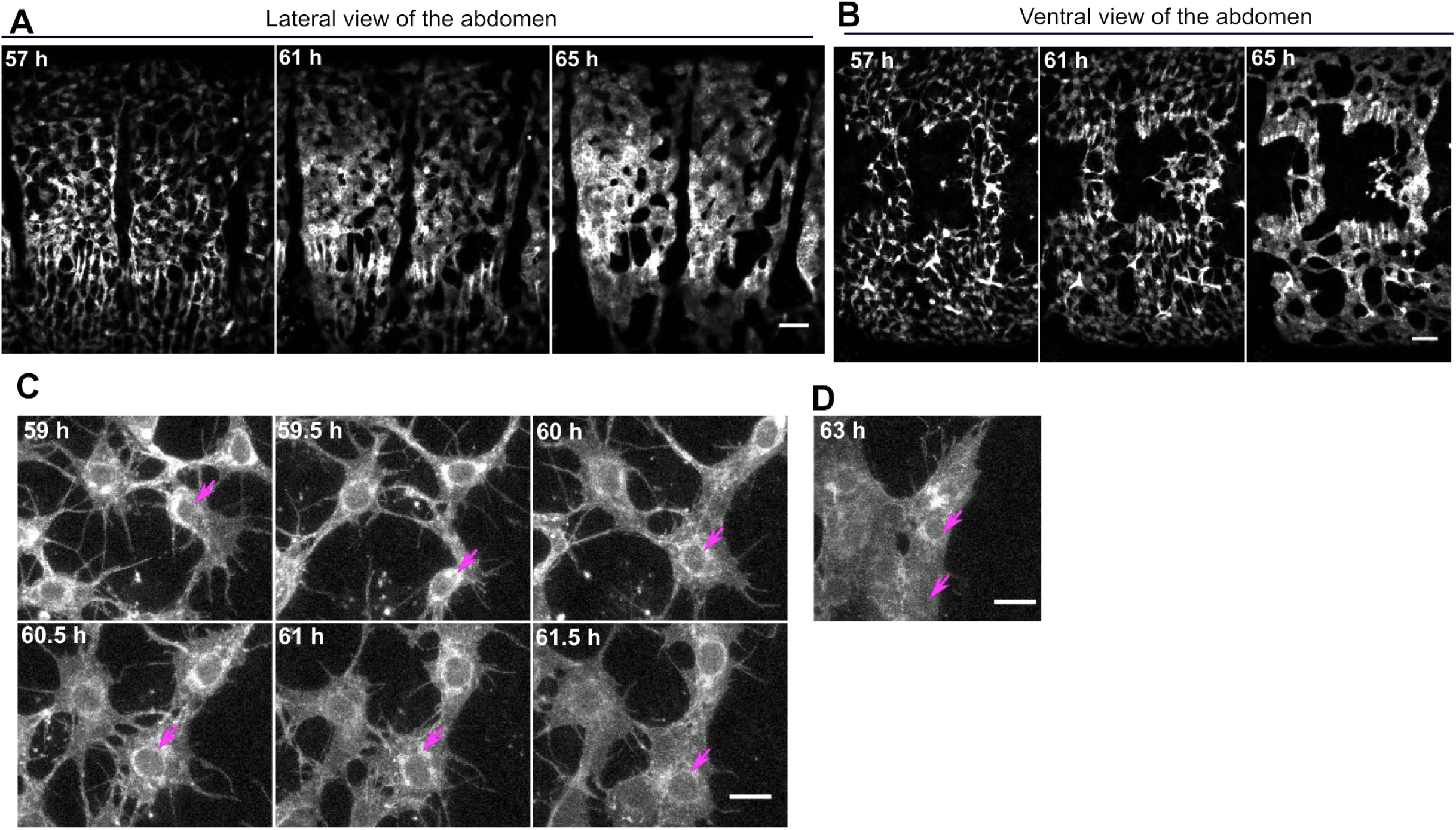
Neighboring AFBp cells adhere to each other to form the monolayer AFB. (A,B) Time-series image stills show the establishment of the monolayer architecture of the AFB from lateral (A) and ventral (B) views of the abdomen. (C,D) Single-cell resolution image stills of AFBp cells during the sheet formation. The cell indicated by arrows underwent flattening (C, 60 h), and the retraction of protrusions (C, 59-61.5 h). At 63 h, AFBp cells had adhered to each other, resulting in the characteristic monolayer arrangement (D; the cell underwent cell division). Scale (μm): 50 (A,B), 10 (C,D).

### Precursor cells may emigrate from the thorax

Our time-lapse imaging suggests that AFBp cells are derived from the thorax or head. To better characterize the site of origin, we attempted to visualize migrating AFBp cells in the thorax and head. While the thick ventral integument of thoracic segments prevented imaging AFBp cells within the thorax, AFBp cells in the head could be imaged in dorsal and frontal views. To observe AFBp cells in the head, we used the *c833-Gal4* and *ppl-4kb-Gal80* combination since *svp[31F09]-Gal4*-based labeling methods failed to visualize AFBp cells in the head for unknown reasons (N = 7 flies with the developed AFB in the abdomen, Fig. S6A). In mature adult flies, the AFB was located beneath the vertex, the frons, and the stem region of the proboscis (Fig. 5A, Fig. S6B-D; Miller, 1950). In the frontal view, AFBp cells seemed to emerge from the border between the head and thorax at approximately 25 h APF, when reared at 29°C (Fig. 5B). They migrated in the anterior direction at the region between the compound eye and proboscis, and a subset of cells entered the proboscis (Fig. 5C, Table 9, Movie 13). Moreover, the displacement length of anterior movements might be slightly larger compared to migration events toward other directions (Table 10). When imaged from the dorsal view, AFBp cells appeared to emerge at the posterior edge of the vertex and exhibited anteriorly biased motility (Movie 14). Taken together, precursors in the head migrated anteriorly, as opposed to abdominal AFBp cells, which moved posteriorly.

**Fig. 5.**
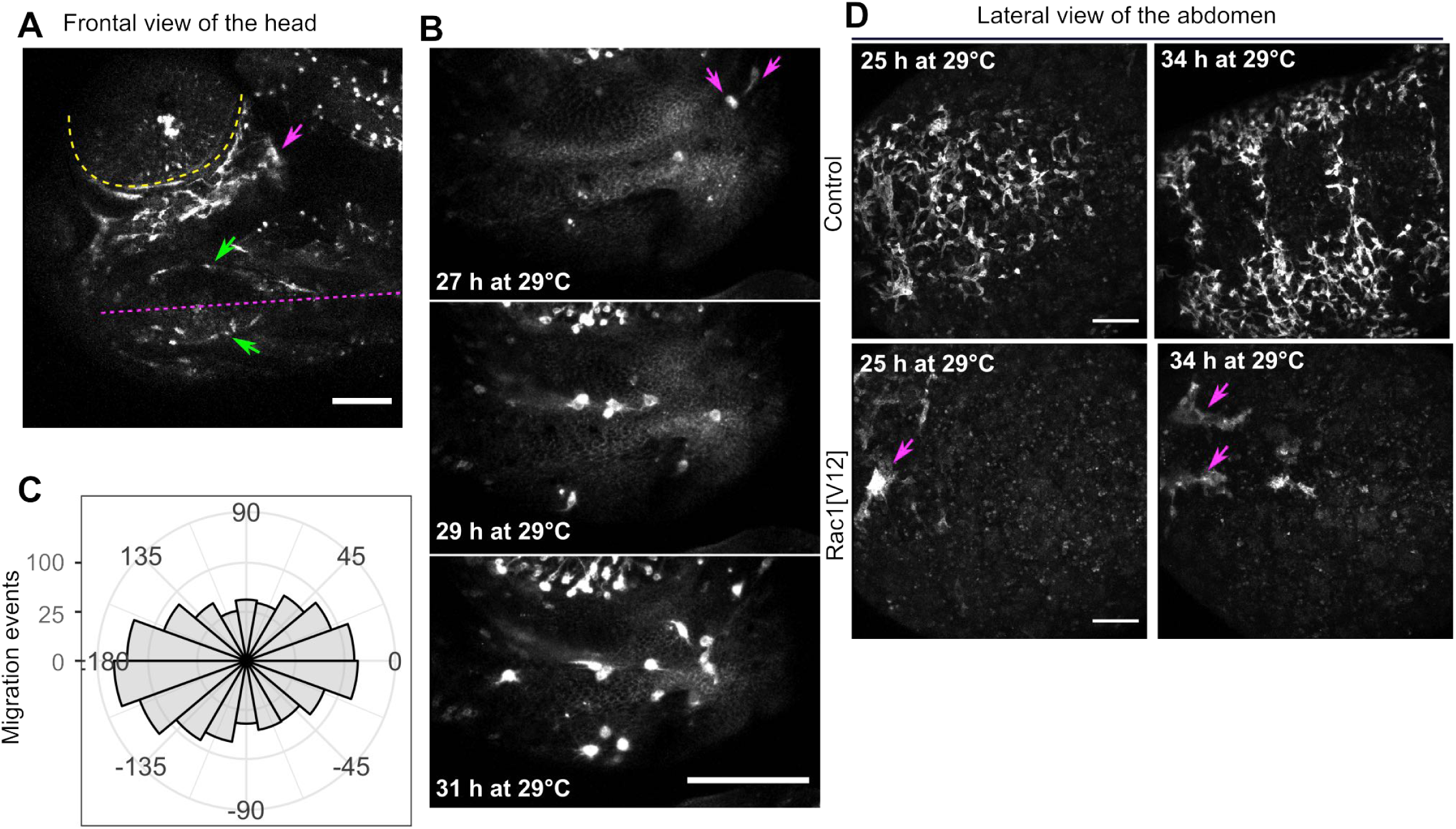
AFBp cells may originate from the thoracic segments. (A-C) AFBp cells in the head migrated in the anterior direction. In the frontal view of the head at 44 h (29°C), AFBp cells were found in regions between the compound eye (yellow dashed line) and the stem of the proboscis at the VML (magenta dashed line). Magenta and green arrows indicate AFBp cells around the compound eye and the proboscis, respectively. (B-C) Time-series image stills of AFBp cell migration (B) and quantification of the orientation of tracked steps at 3 min intervals (C; 1457 steps from 3 female flies). (D) Image stills of the ventral side of the abdomen of female flies show the formation of large cell clumps caused by Rac1[V12] (arrows), possibly by aberrant homotypic adhesions. Note that GFP-positive migrating cells are almost absent in the posterior regions in Rac1[V12] images. Ectopic Rac1 mutants plus GFP expression were induced after puparium formation using the TARGET system (see Materials and Methods). Scale: 100 μm.

These lines of evidence suggest that AFBp cells originate from the thorax; however, there might be a distinct population of abdominal cells that give rise to AFBp cells, and these resident precursors would be concealed by a large number of cells coming from the thorax. To address this possibility, we disrupted the motility of AFBp cells during the early development of the AFB and tested whether hypothetical resident AFBp cells could be detected in the abdomen. We took advantage of mutant variants of Rac1, which acts as a major regulator of cell motility by controlling cytoskeletal dynamics (Luo et al. 1994; Petrie et al., 2015; Pocha et al., 2014). Either dominant-negative or constitutively active forms of Rac1 were expressed in the AFBp cells only after puparium formation to avoid defects before metamorphosis using the TARGET system (McGuire et al., 2004). When the *c833-Gal4* and *ppl-4kb-Gal80* combination was activated by shifting to a restrictive temperature (29°C) after puparium formation, GFP signals within AFBp cells were first detected at 20-25 h APF. At 50-55 h APF, a dominant-negative form of Rac1 strikingly reduced the number of AFBp cells in the abdomen, suggesting Rac1 acts in the migration or proliferation of precursors (Fig. S7, Table 11, Movie 15). When constitutively active Rac1[V12] was ectopically induced, abnormally large cell clumps were seen at the anterior region of the ventral side, suggesting aberrant homotypic cell-cell adhesions between AFBp cells migrating out of the thorax (Fig. 5D, Movie 16). These clumps disappeared by 40 h APF. In the ventral aspect of the abdomen of flies expressing Rac1[V12], a small population of migrating AFBp cells was seen. We speculate that these cells started expressing the marker after the entry into the abdomen and were unlikely to be AFB primordial cells within the abdomen because their emerging sites were inconsistent among the individual flies that we observed (N = 6 flies).

### Continuous cell proliferation and homotypic cell fusion contribute to the establishment of proper tissue mass and multinucleated cells

In newly eclosed adult flies, the abdominal AFB contains approximately 18,000 and 12,000 nuclei in the female and male abdomen, respectively (Johnson and Butterworth, 1985). Achieving the numerous nuclei would require multiple rounds of cell division for each precursor cell, but how and when these cells divide has been unexplored. Our time-lapse imaging captured AFBp cells undergoing cell division during and after migration.

The cell division of migrating AFBp cells involved several distinct steps. Actively migrating cells retracted their prominent protrusions and rounded (Fig. 6A). Concurrently, nuclear-localized fluorescent marker signals spread in area and decreased in intensity, indicating the onset of nuclear membrane breakdown. In these rounded cells, mitotic spindles were observed, as visualized by Clip::GFP, a microtubule marker based on the microtubule-binding domain of human *CLIP170* (Fig. 6B; Stramer et al., 2010). After cytokinesis, the resultant daughter cells recovered their polarized shapes and started to migrate individually (Movie 17). Quantification of the nuclear doubling time of ventral AFBp cells during 25-60 h APF (Fig. 6C, Table 12) revealed their continuous proliferation, perhaps with more rapid nuclear division rates at earlier developmental periods (doubling time for 25-40 h = ∼7.9 h; for 45-60 h = ∼14.6 h). Assuming that the nuclear doubling rate at 15-25 hr APF, when we did not quantify cell proliferation rate due to the paucity of AFBp cells, is the same as that of 25-40 h APF, precursors may undergo approximately four rounds of nuclear division during 15-60 h APF.

**Fig. 6.**
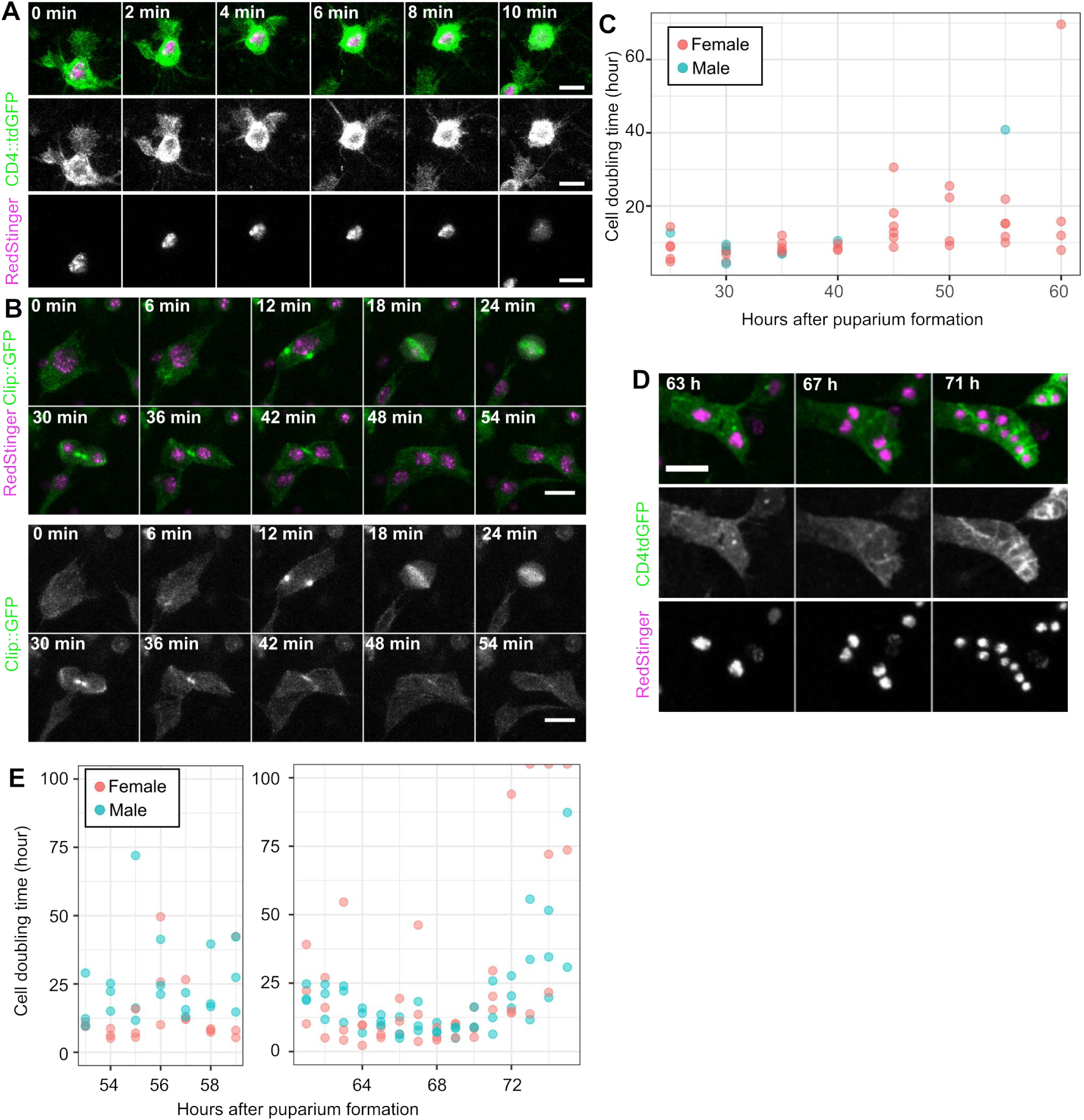
AFBp cells proliferate during and after the migration period. (A) An actively migrating cell retracted protrusions and rounded. Simultaneously, RedStinger (nuclear-targeted dsRed) signal intensities decreased (10min). (B) A mitotic spindle in a spherical AFBp cell was visualized by Clip::GFP. (C) Quantification of nuclear division events of AFBp cells in the ventral aspect of the abdomen from 25-60 h APF. Flies expressing RedStinger were imaged for two hours at 5 min intervals. Each dot indicates the cell doubling time calculated for a single fly (1251 division events in total from 43 flies). These doubling time values are slightly overestimated (see Materials and Methods). (D,E) AFB cells continued dividing after the migration period. Two cells at the tergite formed a lobe-like tissue while undergoing two rounds of cell division, with their total area almost constant, resulting in smaller cells (D). Quantification of the nuclear division rate of AFBp cells in the tergite (E). Nuclei were imaged at 12 min intervals during 53-60 h or 61-80 h APF (184 and 1098 division events from 6 flies, respectively). Each dot indicates a doubling time value calculated from a single fly for a one-hour time window. Data points with greater than 100 hours were excluded. Scale: 10 μm.

Cell division events continued even after AFBp cells were arranged as the monolayer tissue (Fig. 6D, Movie 18). During this post-migration proliferation period, its tissue size was relatively unchanged, but the cell and nuclear sizes of AFB cells were progressively decreased. Adult fat cells showed a rapid decrease in their division rate after 70 h APF, and, by approximately 75 h APF, almost all had ceased their cell cycle progression (Fig. 6E, Table 13). There were 3.3 times as many AFBp cells at 75 h APF as at 61 h APF, implying approximately 1.7 rounds of nuclear division events per cell after 61 h APF, on average.

AFB cells in dipteran species, including *Drosophila melanogaster*, are multinucleated (Day, 1943; Doane, 1960). Multinucleation was also seen in the larval fat body in non-dipteran species (Nakahara, 1918); however, how multinucleated fat body cells are formed has been unclear. We often captured the occurrence of cell-cell fusions between neighboring AFB cells and the resultant multiple nuclei in a single cell from 70 to 85 h APF (Fig. 7, Movie 19). These data argue that homotypic cell-cell fusion between AFB cells contributes to the formation of multiple nuclei found in the imago.

**Fig. 7.**
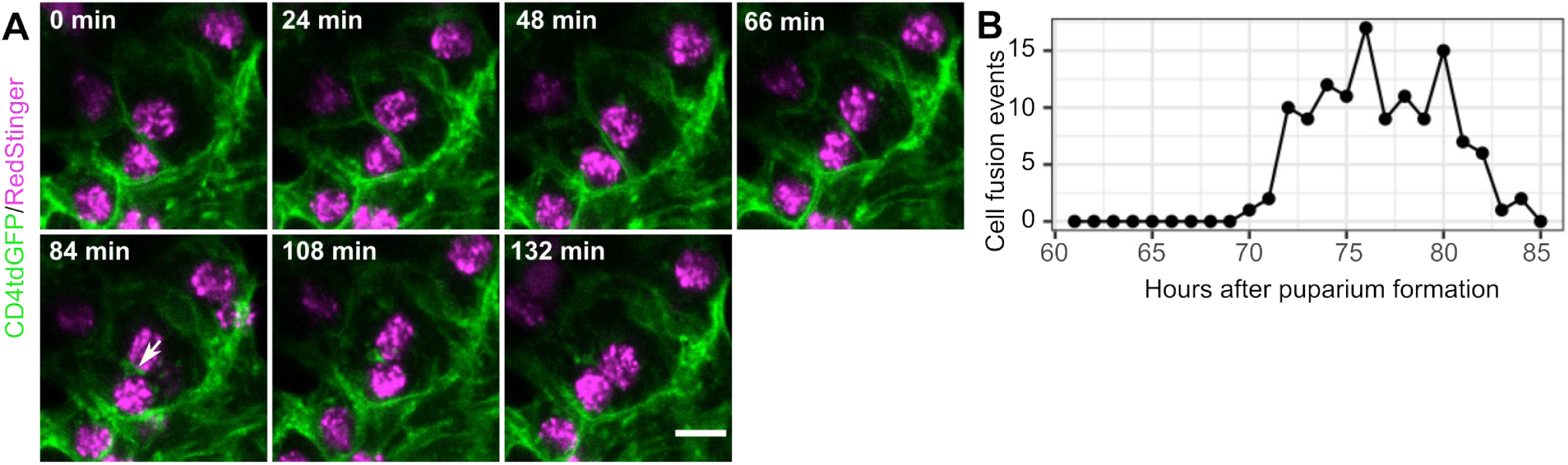
Homotypic cell fusion contributes to multi-nucleated adult fat body cells. (A) Time-lapse imaging, showing the disappearance of cell membranes at a contact site of two neighboring AFB cells in the tergite (arrow), indicating a homotypic cell fusion event. (B) Quantification of cell fusion events from 61-85 h APF at the tergite (N = 122 cell fusion events from 3 flies). Scale: 5 μm.

### Hemocytes are not the primary cellular origin of the adult fat body

Anatomical studies on the development of the AFB in dipteran species suggested that certain types of hemocytes might give rise to adult fat cells (Jones, 1962; Sakurai, 1977; Whitten, 1964). Given that recent studies have shown that *Drosophila* hemocytes show trans-differentiation capacity within hemocyte lineages (Csordás et al., 2021), we examined whether hemocytes directly contribute to the AFB.

First, we imaged hemocytes in the abdomen during mid-metamorphosis (from 24 to 50 h APF) by using *srp.Hemo-Gal4*, a pan-hemocyte Gal4 driver line (Brückner et al., 2004; Evans et al., 2014). There were two types of hemocytes, plasmatocytes and possibly crystal cells (Banerjee et al., 2019). Plasmatocytes exhibited massive intracellular inclusions, some of which contained muscle debris (Fig. S8A,B; Ghosh et al., 2020; Regan et al., 2013). Crystal cells showed no large intracellular inclusions, stronger GFP signals induced by *srp.Hemo-Gal4*, and rounded bulges from the cell membrane at 48 h APF (Fig. S8C). These hemocytes showed distinct cell morphologies and distributions compared to those of AFBp cells (Fig. S8 D-F). In addition, both types of hemocytes in the abdomen showed active migrations with relatively small directional biases during 25-45 h APF, but then their migration activities decreased (Movie 20 and 21). We also examined whether or not the AFB after its formation (72-75 h APF) shows hemocyte-Gal4-driven GFP signals. We observed flies expressing GFP under the control of *srp.Hemo-Gal4* or *pxn-Gal4* (Stramer et al., 2005; N = 3 for each condition) and found no GFP signals in most AFB cells (Fig. S9). These observations indicate that hemocytes are not the primary cellular origin of the AFB.

### Ecdysteroid signaling and Srp are required for adult fat body development

Lastly, we examined the roles of genes that are expected to be crucial for the metamorphic development of the AFB. Ecdysteroids are the central humoral regulator of developmental transitions in insects, including molts and inter-molt events (Thummel, 2001; Truman and Riddiford, 2019). Ecdysteroids are incorporated by target cells and trigger ecdysteroid responses mediated by a heterodimer complex composed of the nuclear receptor EcR and its co-receptor Ultraspiracle (USP; Yao et al., 1993). During metamorphosis, ecdysteroid signaling controls the progression of metamorphosis by promoting breakdown of various larval tissues, triggering remodeling of persistent cells, or promoting differentiation of imaginal cells set aside during embryogenesis (Brown et al., 2006; Ninov et al., 2007). Ecdysteroids are essential in the progression of the metamorphic remodeling of the LFB (Jia et al., 2017; Liu et al., 2013; Rusten et al., 2004).

We examined the roles of ecdysteroid signaling in AFB development by expressing dominant-negative forms of EcR (EcR[DN]) isoforms, which suppress EcR-dependent gene activation by competing for USP with endogenous EcR receptors (Cherbas et al., 2003; Hu et al., 2003). EcR[DN] expression was temporally induced only after puparium formation, similar to the Rac1 mutant experiment (Fig. 5D, Fig. S7). Expression of various EcR isoforms with dominant-negative mutations led to striking reductions in AFB area size in the abdomen when flies were observed at 65-70 h APF (aged at 29°C) and 0-1 day after adult eclosion (Fig. S10, Table 14). Time-lapse imaging revealed that the defects occurred late in the AFB development (Fig. 8A). When imaged 45-50 h APF at 29°C, AFBp cells expressing EcR.A[W650A] or EcR.B1[W650A] in the tergites showed minor defects. During 50-70 h APF, these cells exhibited atrophic changes, and most of them were progressively lost (Movie 22 and 23). These data suggest that ecdysteroid signaling promotes AFB organogenesis or adult fat cell survival through transcriptional activation late in metamorphosis, which may contrast with roles for ecdysone signaling in the induction of histolytic processes in larval fat cells during metamorphosis.

**Fig. 8.**
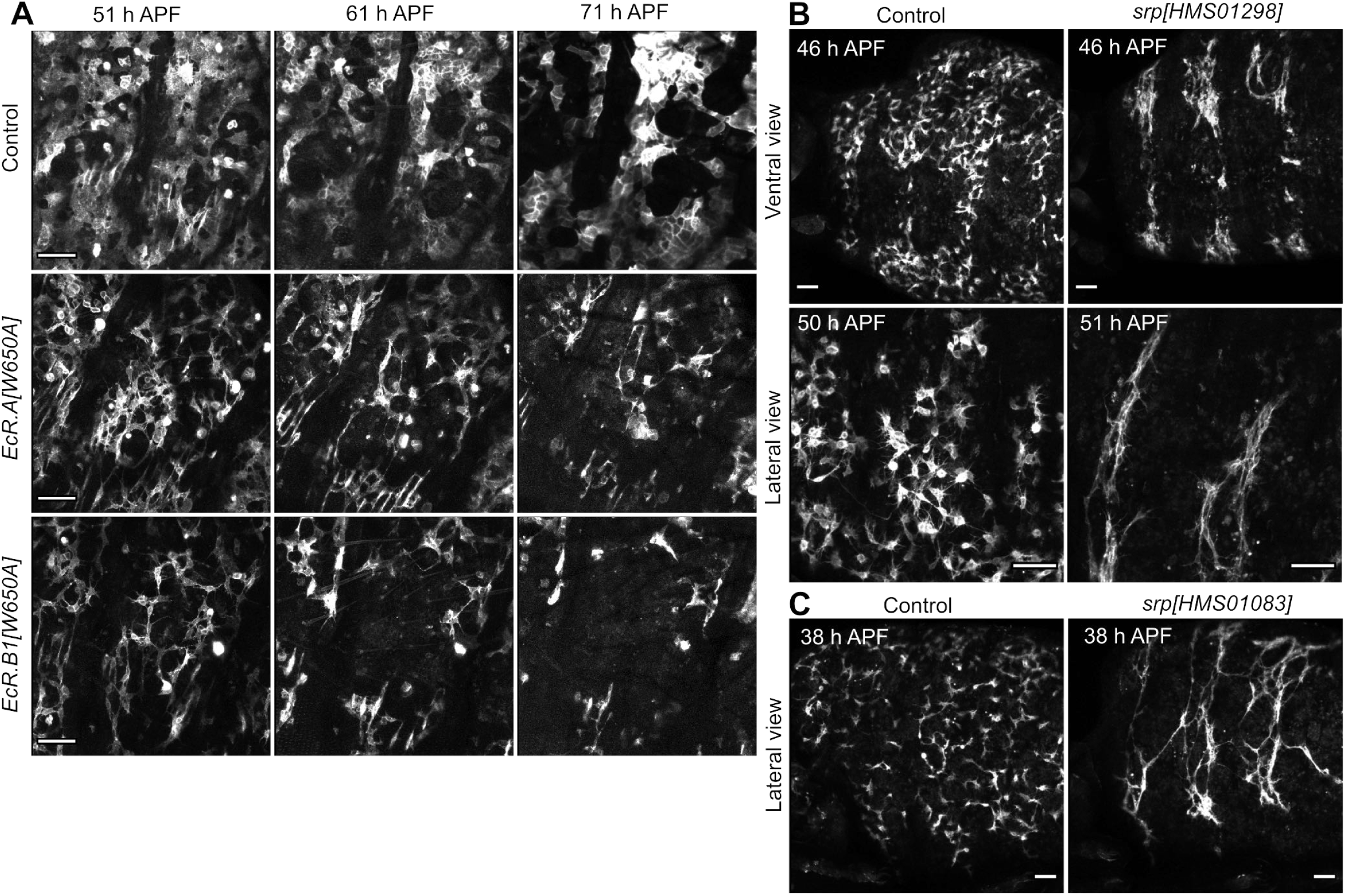
Ecdysteroid signaling and Srp are required for the development of the adult fat body. (A) Ectopic expression of dominant-negative mutants of EcR isoforms disrupted the AFB formation. Ectopic expression was induced after puparium formation, as in Fig. 5. Representative image stills of adult fat cells in the tergite expressing mCherry::CAAX (control), EcR.A[W650A], and EcR.B1[W650A] (dominant-negative mutants). AFBp cells expressing EcR mutants were progressively lost. Nile Red staining also confirmed the significant loss of the AFB in EcR[DN]-expressing mutants but not in EcR[WT]- overexpressing animals (Fig. S10). (B-C) *srp* may be required for the early development of the AFB. Temporal knockdown of *srp* using *srp[HMS01298]* (B) and *srp[HMS01083]* (C) resulted in the formation of cellular assemblies possibly caused by homotypic cell-cell adhesions. Scale: 50 μm.

We also examined the role of the *srp* gene, a GATA-type transcription factor expressed in the earliest stage of the LFB development (Sam et al., 1996; Tremblay et al., 2018). *srp* loss of function results in the absence of larval fat cell precursors by late stage 15, possibly due to precocious cell death (Sam et al., 1996). Moreover, it was reported that forced *srp* expression in the mesodermal lineage resulted in the formation of ectopic larval fat cells, which merged with endogenous fat cells to form an expanded LFB (Hayes et al., 2001). First, we attempted to systemically knockdown *srp* function after puparium formation using *tub-Gal4* and *tub-Gal80[ts]*. We tested five RNAi lines, of which two RNAi lines, *srp[HMS01083]* and *srp[HMS01298]*, caused almost complete loss of the Nile Red-positive AFB at 96 h APF (29°C) with complete penetrance (Fig. S11A; Table 15). These two RNAi constructs do not share target sequences. When driven by *c833-Gal4* and *ppl-4kb-Gal80*, *srp[HMS01298]* resulted in a dramatic loss of AFBp cells at 50 h, when aged at 29°C (Fig. S11B), reminiscent of lost larval fat cells in *srp* mutant embryos. When *srp[HMS01298]* expression was induced later in development (from 24 h APF aged at 17°C onward), migrating AFBp cells frequently showed thread-like cellular assemblies (Fig. 8B, Movies 24 and 25), suggesting aberrant homotypic cell-cell adhesions among AFBp cells. An independent knockdown induced by *srp[HMS01083]* resulted in web-like assemblies of AFBp cells (Fig. 8C, FigS11B, Movies 26 and 27). These similar defects in AFB assembly, likely due to aberrant adhesions among AFBp cells, suggest that the aberrant AFB developmental phenotypes are in fact due to Srp depletion and not off-target effects.

## Discussion

Early anatomical studies in dipteran species revealed that the AFB newly develops by adult eclosion (Evans, 1935; Pérez, 1910; Robertson, 1936; Wigglesworth, 1949); however, how the AFB develops during metamorphosis has remained unclear. Here, we identified highly migratory precursor cells and characterized the cellular dynamics underlying AFB development at the single-cell resolution. Precursor cells underwent a long-distance migration, continuous proliferation, and structural transformation to form the monolayer. These cellular and tissue-level events appeared to be temporally and regionally controlled (Fig. 9). AFBp cells were assembled into the AFB and started lipid deposition by approximately 65 h APF, consistent with reports of the accumulation of lipid droplets in the AFB during metamorphosis in dipteran species other than the fruit fly (Evans, 1967; Wiessmann, 1962). Thus, AFB development in flies will provide a unique experimental platform in which most of the developmental events of an adipose tissue can be observed in vivo.

**Fig. 9.**
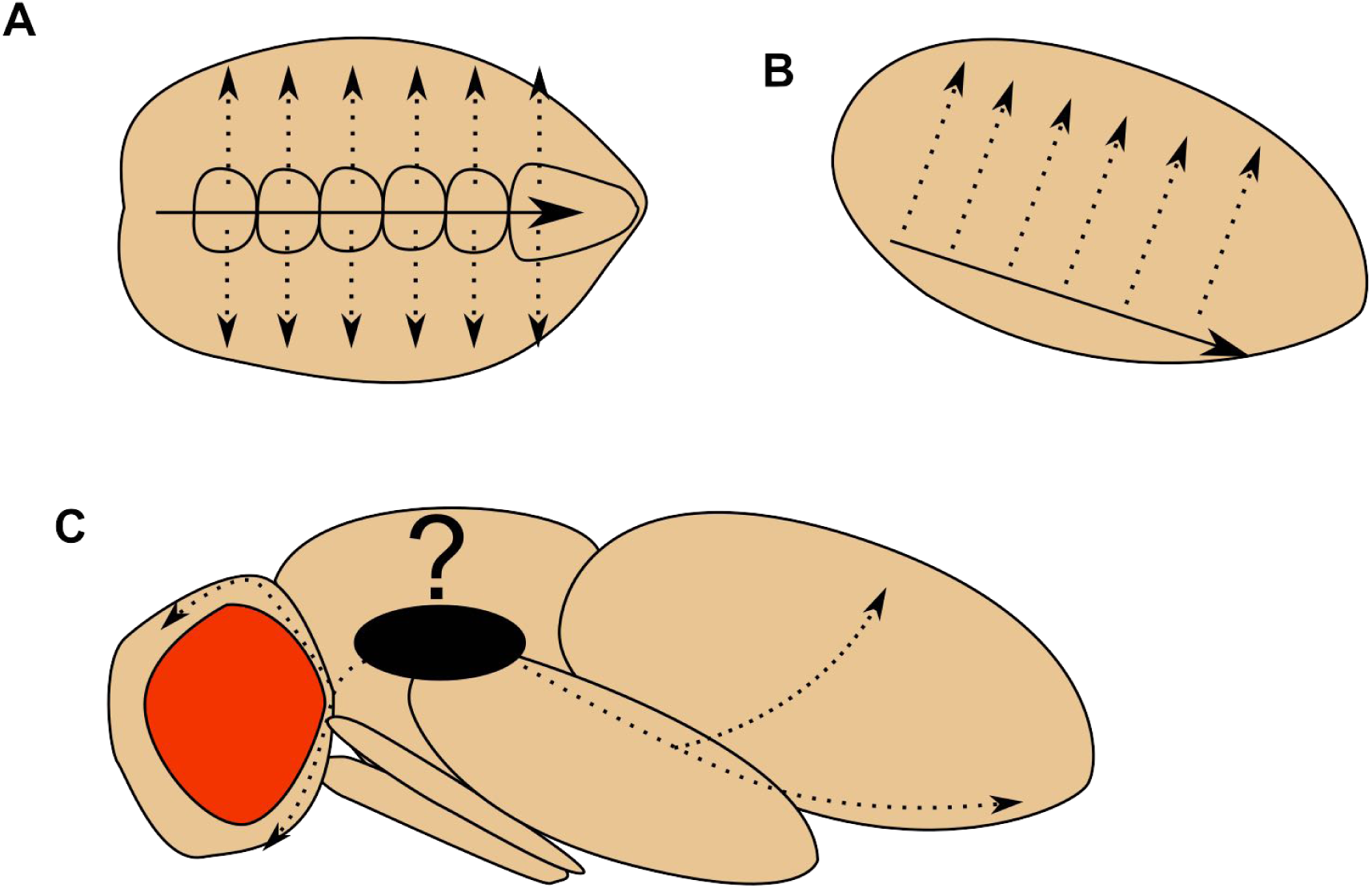
AFBp cells undergo a long journey to disperse across the whole body. Schematic illustrations of AFBp migration patterns in the abdomen (A,B) and the site of the origin of the AFB (C). In the ventral abdomen, AFBp cells emerged at the anterior end (15 h APF) and migrated predominantly in the ventral posterior direction at first (15-30 h APF; arrows in A,B). Then, a subpopulation of these cells started to migrate dorsally (30 h APF; dashed arrows in A,B) and reached the dorsal midline (50 h APF). While our data support the notion that AFBp cells originate from thoracic segments, the detailed site of origin is still unclear (C).

We established genetic tools to highlight the developing AFB during metamorphosis. First, we employed a 4-kb cis-element in *ppl-Gal4* to induce *Gal80* in the LFB (Buch et al., 2008; Zinke et al., 1999). The *ppl-4kb-Gal80* system might be helpful for labeling other cell types when used with Gal4 drivers expressed both in cells of interest and the LFB. Second, we found that two *svp-Gal4* strains in Gal4 libraries (Jenett et al., 2012; Kvon et al., 2014) showed a transient expression in an AFB lineage at an early stage. While our AFB Gal4 tools provided important insights into cellular mechanisms, their utility was limited due to undesired expressions in neighboring tissues and the difficulty in precisely timed control of Gal4 function (see Materials and Methods). Nonetheless, the basic strategy will be useful for further refinements to genetically dissect mechanisms underlying AFB development.

Our data argue that AFBp cells emigrate from the thorax and undergo a long-distance journey to disperse across the body. The broad distribution is likely critical for circulating energy metabolites and signaling molecules to respond quickly environmental challenges and avoid adverse effects due to locally accumulated metabolites. The migration-based distribution strategy appears contrast with those of other mesodermal organs. Larval fat cell precursors differentiate in a metameric manner in the embryo, followed by cell proliferation and assembly into the LFB (Azpiaz et al., 1996; Hoshizaki et al., 1994; Miller et al., 2002). Larval and adult muscles, another class of mesodermal tissue undergoing metamorphic tissue remodeling, are also locally generated during embryogenesis and metamorphosis, respectively (Bate et al., 1991; Baylies, et al., 1998; Currie and Bate., 1991; Gunage et al., 2017). Thus, on-site differentiation is likely to be the canonical mechanism for the broad distribution of mesodermal organs in the fly.

If AFBp cells were generated in central segments, their adequate distribution across the whole body would be problematic. Since the AFB is abundant in the abdomen of the imago, especially in the tergite, most precursors must migrate posteriorly to reside in the abdomen, and a remaining minor population can migrate into the head. Moreover, the abdominal population must preferentially migrate dorsally to reach the tergites, with a subset of the population remaining in the pleurite and sternite. The directional switching around the abdominal spiracles may help to prevent their excessive dorsal colonization. While we did not address the molecular mechanisms underlying the allocation, these mechanisms could be intrinsic, extrinsic, or a combination of both. Differential gene expression profiles between the head and abdominal AFBs in the adult fly have been reported (Fujii and Amrein, 2002). If the distinction in gene expression profiles is established during early differentiation, intrinsic factors could direct different populations toward different destinations. Such an early genetic separation scenario might be consistent with the fact that *svp[31F09]-Gal4* plus *Ay-Gal4*, a potent Gal4 driver for the abdominal AFB, failed to be expressed in the head AFB.

Extrinsic factors provided by other tissues and organs may control AFBp cell dynamics. The migration area for AFBp cells around the ventral midline progressively narrowed during 25-35 h APF, and a significant subset of AFBp cells started to migrate dorsally at 30 h APF. During the same developmental period, the ventral nest histoblasts, which form imaginal abdominal epidermis, continued to expand ventrally, and replaced the larval epidermal cells by 32 h APF (Movie 28; Ninov et al., 2007; Roseland and Schneiderman, 1979). In addition, adult lateral muscles in the pleurite start to elongate along the dorsoventral axis at 31 h APF (Currie and Bate, 1991). These remodeling events might shape AFBp migration patterns by physically constraining their migration space or providing scaffolds and attractants for the directed migration. Another candidate tissue controlling AFBp cell migration may be oenocytes, which predominantly reside in the tergites and are functionally connected with the AFB in lipid circulation (Bousquet et al., 2012; Chatterjee et al., 2014; Koch, 1945). Additionally, oenocytes intermingle with fat cells to assemble into the fat body in various insect species (Makki et al., 2014). These facts raise the possibility that oenocytes secrete humoral factors that attract AFBp cells. While developmental mechanisms of adult oenocytes have been still unclear, our time-lapse imaging revealed that developing adult oenocytes appeared to emerge by 35 h APF (Movie 29 and 30), almost concomitant with the onset of the dorsal migration of ventral AFBp cells.

Continuous proliferation during the migration period appears to be a prominent feature of AFBp cell dynamics and is inherently necessary to build up the AFB in the limited time frame for metamorphic transformation. Cell proliferation during cell migration has been reported in various cell types, including neural crest cells, neuroblasts, and germ cells in mammals (Cantú et al., 2016; Ridenour et al., 2014; Zhang et al., 2007). The proliferation during migration may be advantageous because migrating cells could sense environments along their paths and at destinations. For instance, starved fly larvae can progress into metamorphosis when they have passed the critical timing, resulting in small adult flies (Beadle et al., 1938). AFBp cells in these small flies could sense changes in body size and match their tissue mass by modulating their proliferation or cell-size growth.

Where AFBp cells originate has a long-standing question. Several studies in dipteran species suggest the possibility that a subset of hemocytes transdifferentiate into fat cells and form the AFB (Jones, 1962; Sakurai, 1977; Whitten, 1964). Our data imply that hemocytes do not provide a significant direct contribution to the AFB. It might be challenging to distinguish polarized and motile AFBp cells from hemocytes, which are often highly migratory and load various types of debris, including lipid-containing structures (Ghosh et al., 2020; Jones, 1962). Quiescent larval cells associated with the epidermis in abdominal segments have also been proposed as progenitors in histological and histochemical studies (Koch, 1945; Wiesmann, 1962; Hoshizaki, 1995). Our time-lapse imaging argues that AFBp cells originate from central segments but not abdominal segments. Adepithelial cells associated with wing and eye-antennal imaginal discs would be the sole previously proposed candidate cellular origins that are consistent with our observations (Hoshizaki et al., 1995); however, a previous wing imaginal disc transplantation study did not report the identification of adult fat cells (Lawrence and Brower, 1982). Thus, the specific site and origin of the AFBp cells is still unclear.

De novo formation of the AFB during metamorphosis has only been reported in limited species, including derived dipteran species (reviewed in Dean et al., 1985; Trager, 1937). In other holometabolous insect species, it is widely accepted that the embryonically developed fat body operates throughout the life cycle, with a large-scale reorganization of intracellular structures during metamorphosis (Dortland and Esch, 1979; Haunerland and Shirk, 1995; Larsen, 1976; Oertel, 1930). Therefore, the de novo formation of the adipose tissue during metamorphosis seems to have evolved in derived species. Our results suggest that AFB development may involve common transcription programs used in LFB development, including Srp and Svp. Larval and adult fat cells may branch from a common ancestral cell lineage, which could make up the embryonic fat body in basal species. The diversification in differentiation programs might involve the modulation of responses to humoral factors (Butterworth 1972), including ecdysteroids. In a few Lepidopteran species, regionally and biochemically distinguished embryonically generated fat body tissues take contrasting fates during metamorphosis (Haunerland et al., 1990; Shirk and Malone, 1989; Wang and Haunerland, 1992). While a population of larval fat cells in these species is destined to be degraded during pupal stages, another subset persists into the adult. The elucidation of molecular mechanisms underlying the regional difference might provide a clue to understand how postembryonic AFB formation has evolved.

## Materials and Methods

### Fly strains and husbandry

Flies were raised and maintained at 25°C on standard cornmeal food in plastic vials unless otherwise specified (Watanabe, Furumizo et al., 2017). Fly strains were obtained from the Bloomington Drosophila Stock Center, the KYOTO stock center, the Vienna Drosophila Resource Center, and published resources. Fly strains and plasmids used in this study are summarized in Table 16. Exact genotypes are summarized in Table 17.

Precisely staged flies for time-lapse imaging were obtained by collecting white prepupae on the wall of plastic vials using wet paintbrushes at one-hour intervals (defined as 1 h after puparium formation: APF). The flies were stored for further development in a humid incubator. At the time of live-imaging and histochemical staining, flies were placed on double-sided adhesive tape on glass slides, and their puparium was carefully removed using forceps.

### Genetic tools for Gal4 expression in the adult fat body and oenocytes

For searching for Gal4 stocks that can visualize AFB development, commonly used fat body Gal4 lines were first examined for expression of GFP markers in the LFB and AFB of young adult flies. Gal4 driver lines we tested were *Cg-Gal4*, *r4-Gal4*, *c833-Gal4*, *ppl-Gal4*, and *Lsp2-Gal4* (Asha, et al., 2003; Buch, et al., 2008; Lee and Park, 2004; Hrdlicka, et al, 2002). These Gal4 lines showed strong expression in the persistent LFB. This persistent expression prevented delineating AFB development, even though *c833-Gal4* was expressed in the AFB lineage as revealed by Kaede, a photoconvertible protein emitting green or red fluorescence before and after photoconversion, respectively (Fig. S1C; Ando et al., 2002; Tsuyama et al., 2017). Photoconversion was performed using a macro zoom microscope (MVX10, Olympus) with a mercury lamp and a band-pass filter transmitting ultraviolet wavelength light (U-MWU2, Olympus).

Gal80 lines under the control of cis-elements of fat-body-related genes were established (Fig. S1D). The regulatory cis-element sequences were selected based on genomic positions of genes of interest and neighboring genes using Flybase (Thurmond, et al., 2018) and amplified by PCR reactions (KOD FX, KFX1-1, Toyobo) from genomic DNA or a plasmid containing a cis-element of the *ppl* gene (pCK1, Buch et al., 2008; Table 1). Primers used to amplify fragments are summarized in Table 1. Amplified DNA fragments were subcloned into entry vectors (pENTR/D-TOPO, K240020, ThermoFisher; or pCR8/GW/TOPO, K 250020, ThermoFisher) and then were inserted into pBP-Gal80Uw-6 (#26236, Addgene) via an L-R reaction with GATEWAY LR clonase II plus enzyme mix (12538120, Thermo Fisher). pBP-Gal80Uw-6 was designed to establish constructs carrying Gal80 under the control of regulatory cis-elements of interest (Pfeiffer et al., 2010). Fly strains carrying fat-body Gal80 strains were established by a commercially available injection service company (Best Gene Inc). The landing site used was the attP2 site. Among the Gal80 stocks generated, *ppl-4kb-Gal80* was utilized with *c833-Gal4* to image AFBp cells. The DNA sequence used as the cis-element of our *ppl-4kb-Gal80* strain contains the *ppl* coding sequence and approximately 3kb of flanking sequences (Buch et al., 2008). *ppl-4kb-Gal80* also suppressed Gal4-dependent marker expression in hemocytes driven by *pxn-Gal4* at the late third instar and white prepupa stage (Fig. S3).

In our search for strains expressing Gal4 early in the AFB lineage, the G-TRACE system was employed to avoid overlooking strains with transient Gal4 expression (Evans et al., 2009). In G-TRACE, a nuclear-targeted GFP marker (Stinger) under the control of a ubiquitin promoter represents lineage signals. In the absence of Gal4 activity, the ubiquitin promoter and the open reading frame of Stinger are separated by a transcriptional termination signal within an FRT cassette. After Gal4 expression, activation of *UAS-FLP* causes the irreversible removal of the FRT cassette, resulting in persistent Stinger expression as a memorized lineage signal. In addition, *UAS-RedStinger* was employed to monitor real-time Gal4 function. Gal4 lines related to *serpent* (*srp*) and *seven-up* (*svp*) were tested for G-TRACE signals. *svp[31F09]-Gal4* and *svp[31H02]-Gal4* showed GFP signals (lineage) but not RFP signals in the AFB in young adult flies (Fig. S2F), suggesting that these lines expressed Gal4 in precursor cells in the early development of the AFB but did not maintain Gal4 expression through to adult eclosion. *svp[31F09]-Gal4* was used to image AFBp cells combined with *Ay-Gal4* to sustain Gal4 activity in the AFB lineage (Fig. S2G; Ito et al., 1997).

To visualize early developmental events of adult oenocytes, we first tested *PromE-Gal4*, a widely used oenocyte-specific Gal4 line (Bousquet et al., 2012) and *svp[32C04]-Gal4*. We found that *svp[32C04]-Gal4* was expressed in adult oenocytes in newly eclosed adult flies in our fat-body Gal4 search. These Gal4 drivers began to be expressed in oenocytes later in metamorphosis. Next, we examined *Gal4* lines associated with genes required for early differentiation and delamination of larval oenocytes (Brodu et al., 2004). *vvl[17C04]* and *slm[LP39]* strains enabled relatively specific labeling of adult oenocytes from 25 and 30 h APF onward, respectively.

### Microscopic observation

Flies were imaged using Nikon-C1 confocal laser scanning microscopic systems equipped with an inverted microscope (Eclipse Ti, Nikon) or an upright microscope (Eclipse E800, Nikon). Objectives used were 10x (NA 0.3 or 0.4), 20x (NA 0.75), 40x (NA 1.3, oil-immersion), and 60x (NA 1.4, oil-immersion).

For imaging at a single time point without fixation, whole flies or the abdomens dissected away from the thorax were imaged. Whole live flies were placed on glass-bottom dishes or glass slides and then imaged with an inverted microscope or upright microscope, respectively. The separated abdomens were mounted in 50% glycerol in phosphate-buffered saline, with a piece of plastic tape as a spacer to avoid crushing the abdomen.

For time-lapse imaging, flies were placed on glass-bottom dishes with their puparium removed and imaged using an inverted confocal microscopy. For observations with oil-immersion objectives, flies were placed on a thin layer of halocarbon oil 27 (H8773, Sigma-Aldrich) used as the imaging medium. For ventral view imaging, legs and wings were displaced by using forceps as these structures conceal the abdominal surface. A humid environment was maintained by placing a sheet of paper soaked with water in dishes and using a humidifier in the imaging room. For imaging with long-time courses (12-24 hours), flies sometimes underwent drying, especially in their extremities (i.e., wings and legs); however, the onset timings of AFB developmental events in those flies were not strikingly affected when compared with flies aged in a humid incubator, with their puparium intact. We confirmed that imaged flies were able to proceed to the mature pharate adult stage after observation. For most flies observed for time-lapse imaging, their sex was determined by external genital structures at the pharate adult stage or the male-specific genitalia rotation (Inatomi et al., 2019), and noted in Table 17. In both sexes, the dynamics of AFBp cells were similar. While some aspects of AFB development seemed to differ depending on sex (Fig. 6E and Fig. S5F), more detailed studies will be needed to clarify quantitative differences between sexes.

For ectopic expression or knockdown experiments, the TARGET system was employed, where *tub-Gal80[ts]* is used for timed control of Gal4 activation (McGuire et al., 2004). The Gal80[ts] protein was a temperature-sensitive mutant of Gal80, which inhibits Gal4 under permissive temperatures. After crossing, vials were maintained at a permissive temperature (19°C) to inhibit Gal4 in embryonic and larval stages, and prepupae were collected over six-hour periods (0-6 h APF at 19°C). These flies were aged at 19°C until stages specified in figure legends, and then transferred to a restrictive temperature (29°C). *c833*-*Gal4* plus *ppl-4kb-Gal80* was employed as a Gal4 driver. *svp[31F09]-Gal4* plus *Ay-Gal4* based labeling methods were not applicable with Gal80[ts] because *UAS-CD4::tdGFP* expression in AFBp cells was undetectable even when flies were aged at 29°C throughout their development. The partially remaining inhibitory function of Gal80[ts] at 29°C may decrease FLP levels driven by *svp[31F09]-Gal4* and suppress the FRT cassette removal required for the activation of *Ay-Gal4* (Fig. S2G). Female flies were selected for knockdown or ectopic expression experiments because strong auto-fluorescence in the abdomen of male flies aged at 29°C often interfered with visualizing AFBp cells with fluorescent markers.

For imaging AFBp cells in the head, the *c833-Gal4* plus *ppl-4kb-Gal80* combination was used since *svp[31F09]-Gal4*-based labeling methods failed to visualize AFBp cells in the head for unknown reasons (N = 7 flies with the fully developed AFB in the abdomen, Fig. S6A). *tub-Gal80[ts]* was employed to suppress *c833-Gal4* expression in the larval salivary gland during embryonic and larval stages. Without timed induction with *tub-Gal80[ts],* AFBp cells in the head were often concealed by large floating debris with strong fluorescent signals derived from the histolyzed salivary gland (Farkas and Mechler, 2000).

For assessing the rate of AFB development defects in large numbers of wildtype strains that were relatively recently established as inbred lines, *Drosophila* Genome Reference Panel (DGRP) strains were used (Huang, Massouras, et al., 2014). Thirty-nine strains that belong to the core forty strains were first tested by using stereomicroscopic observations without dissection (MS5, Leica), and flies that appeared to display AFB defects and were hard to judge were further examined by subsequent dissection (Table 4). Accuracy of the observation procedure above was assessed as follows. *Canton-S* female flies (7-8 days after adult eclosion; N = 134) were first examined using a stereomicroscope without dissection by taking advantage of the fact that adult flies with defects in AFB development tend to display the deflated, transplanted abdomen. Subsequently, these flies were dissected as described in “Nile Red staining and immunostaining” and examined whether the AFB was developed by observing them in bright-filed view. Without dissection, most *Canton-S* flies were correctly judged whether they developed the AFB or not (N = 87 and 19 flies with the fully developed or underdeveloped AFB, respectively). Some *Canton-S* flies were hard to judge without dissection, and subsequent observations of dissected flies showed that most of these flies exhibited defects in AFB development (N = 21/28).

### Migration quantification

Digital images were processed with a control application for Nikon C-1 confocal systems (EZ-C1, Nikon) and the Fiji software (Rueden et al., 2017; Schindelin et al., 2012). For cell migration tracking, the TrackMate plugin in Fiji was used (Tinevez et al., 2017). Z-projected image sequences were obtained at 1-3 min intervals and analyzed. Positions in the x-y coordinates of randomly selected cells or all of the cells within a ROI were tracked. When possible, the positions of each cell were automatically recorded using the semi-automatic tracking function in TrackMate. When tracked cells were too close to neighboring cells, locations were manually recorded to avoid generating erroneous links. AFBp cells that underwent cell division were excluded from our analysis. Tracked positions in the x-y coordinates were imported into R software (R core team, 2021) to calculate the displacement length and angle value (from -180° to 180°; anterior direction: ±180°, posterior direction: 0°, dorsal direction: 90°, ventral direction: -90°) for each step. Steps were classified into four directional categories with equal angles (i.e., 90°) to address whether migration parameters differed depending on particular directions. In presenting migration angle distributions using rose diagrams, migration steps smaller than 1 μm/min were excluded, since movements with small displacement values might be due to tissue movements or caused by imprecise manual recordings of motionless cells. Our quantification was not largely affected when all recorded steps were included.

### Proliferation and cell fusion quantification

For quantifying the nuclear doubling time of AFBp cells, flies expressing Stinger or RedStinger induced by *svp[31F09]-Gal4 plus Ay-Gal4* were imaged. Nuclear division events in z-projected image sequences were manually counted by taking advantage of reduced signal intensities of nuclear markers, possibly due to nuclear envelope breakdown. Total nuclear numbers were automatically quantified by thresholding images and counting extracted masks using Fiji. For calculating nuclear doubling time in each time window, total nuclear numbers were divided by time frame numbers, multiplied by imaging hours, then divided by the number of observed division events.

For quantification of nuclear division rates in the ventral abdomen from 25 h through 65 h APF, AFBp cells in A3-A6 segments were observed at 5 min intervals for 115 min with a 20x dry objective (NA 0.75; N = 43 flies in total). Oil-immersion objectives were not used for ventral view imaging as imaged flies often died after imaging. The damage might be due to halocarbon oil penetrating flies through narrow slits on their surface caused by forceps used for displacing their legs and wings. Due to the limited spatial resolution of the 20x objective we used, precise recordings of cell numbers were often challenging in areas with a dense population of AFBp cells. We manually analyzed data for 25 h and 40 h APF when AFBp cells were sparse and dense, respectively, and found that unadjusted values were slightly overestimated by 3.6± 3.9% and 6.4±1.3% (mean±SD; N = 6 and 4, respectively) for 25 h and 40 h, respectively.

For measuring the nuclear division rate in the tergite, flies were collected and imaged with a 60x objective at 12 min intervals during 53-60 h or 61-81 h APF (N = 6 flies for each period).

For quantifying cell fusion events in the tergite, the AFB of flies expressing RedStinger and CD4::tdGFP was imaged using a 40x objective during 61-85 h APF at 6 min intervals (N= 3 flies). After identifying multi-nucleated adult fat cells within the field of view in the last frame, fusion events were manually and retrospectively detected in the image sequences. Identified fusion events in our quantification were partial fusion events even in the field of views due to too strong GFP signals obscuring cell-ell boundaries, and imprecise z-projection matching, and large positional changes of cells caused by muscle contractions.

### Nile Red staining and immunostaining

Nile Red was dissolved in acetone at 1 mg/ml (stock solution, stored at 4 °C) and used to stain neutral lipids at 1 μg/ml (144-0811, WAKO). Hoechst 33342 was used to counterstain nuclei at 100 ng/ml (H 3570, Molecular probes). Flies were dissected by using insect pins (Minutein Pins, 26002-10, Fine Science Tools) and micro scissors on sylgard-based dissection stages (SYLGARD 184 Silicone Elastomer Kit, Dow Corning). The persistent larval fat cells were blown out by pipetting. Specimens were then fixed with 3.7% formaldehyde in phosphate-buffered saline for 15-30 min at 25°C prior to Nile Red and Hoechst staining. If necessary, an antibody against GFP (rabbit polyclonal IgG, A11122, Molecular Probes) and a secondary antibody against rabbit IgG conjugated with Alexa 488 (A-11034, ThermoFisher) was used for immunostaining at a dilution of 1:300 and 1:1000, respectively, to augment GFP marker signals. Fixed specimens were mounted in 50% glycerol in PBS with a piece of plastic tape as a spacer to avoid crushing the specimen.

### Statistical analysis and image processing

Statistical analyses were performed using R. The R codes used in this study are available upon request. Figure panels were constructed using the Inkscape software (Inkscape Project, https://inkscape.org/).

To prepare temporal-color coded images, the Temporal-Color Code function in Fiji was used (https://imagej.net/Temporal-Color_Code). Migration angle distributions were illustrated using rose diagrams, which were plotted as bar charts with polar coordinates using the ggplot2 package in R (Wickham, 2016). For time-lapse imaging using pharate adult flies after the onset of muscle contraction, shifts in the x-y coordinates caused by their motions were automatically matched using the Linear Stack Alignment with SIFT function in Fiji (Lowe, 2004).

In estimating the probability of each strain showing a disrupted or absent AFB in various genetic backgrounds (Table 2, 3), hierarchical Bayesian models were used to incorporate statistical shrinkage (Efron and Morris, 1977). We assume that the AFB defect probability of each strain (*θ_i_*) is distributed as a beta distribution as follows;

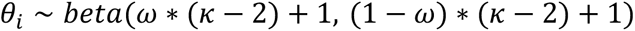

where *ω* and *κ* are the mode and concentration of a beta distribution, respectively.

These hyperparameters were assumed to be distributed as follows.

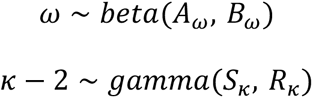

Distributions of lower-level parameters (*θ_i_*) and hyperparameters were estimated using a Jags script for Bayesian binomial inference described in Kruschke (2015). Hyperpriors with weak information were used (dbeta(1, 1) and dgamma(1.105125, 0.1051249)). Maximum a posteriori estimation values and equal-tailed intervals (ETI) of posterior distributions were represented as summary statistics.

To examine the relationship between the migration direction of AFBp cells and the position along the dorsoventral axis (Fig. S5C), we fitted the general additive model (GAM) as

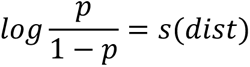

where *p* denotes the probability that observed migration steps are toward the dorsal direction (i.e., 45°-135°), *s*() represents a smooth function, and *dist* is the distance between cellular positions and the VML. The mgcv package in R was used for fitting GAM models to data (Wood, 2017).

## Acknowledgements

Fly stocks and genomic datasets were provided by the Kyoto Stock Center, the Bloomington Stock Center, the Vienna Drosophila Resource Center (VDRC, www.vdrc.at), the TRiP at Harvard Medical School (Grant: National Institutes of Health, Office of the Director R24 OD030002), the Drosophila Genomics Resource Center, FlyBase, P. Martin, A. Gould, C. Han, S. Hayashi, M. Miura, K. Emoto, and M. Pankratz. We also thank J. Hejna for polishing the manuscript; and K. Oki, M. Futamata, and H. Imai for technical assistance.

## Author contributions

T. Tsuyama, K. Shimono and T. Uemura conceived the project. T. Tsuyama, H. Komai, and Y. Hayashi performed experiments. T. Tsuyama performed image and statistical analysis. T. Tsuyama and T. Uemura wrote the paper, incorporating suggestions from all authors.

## Competing interests

The authors declare no competing financial interests.

## Funding

This work was funded by grants from AMED-CREST (JP18gm1110001 to T. Uemura) and Japan Society for the Promotion of Science (08J05328 to T. Tsuyama and 10J06191 to K. Shimono).

## Figure legends

**Fig. S1.**
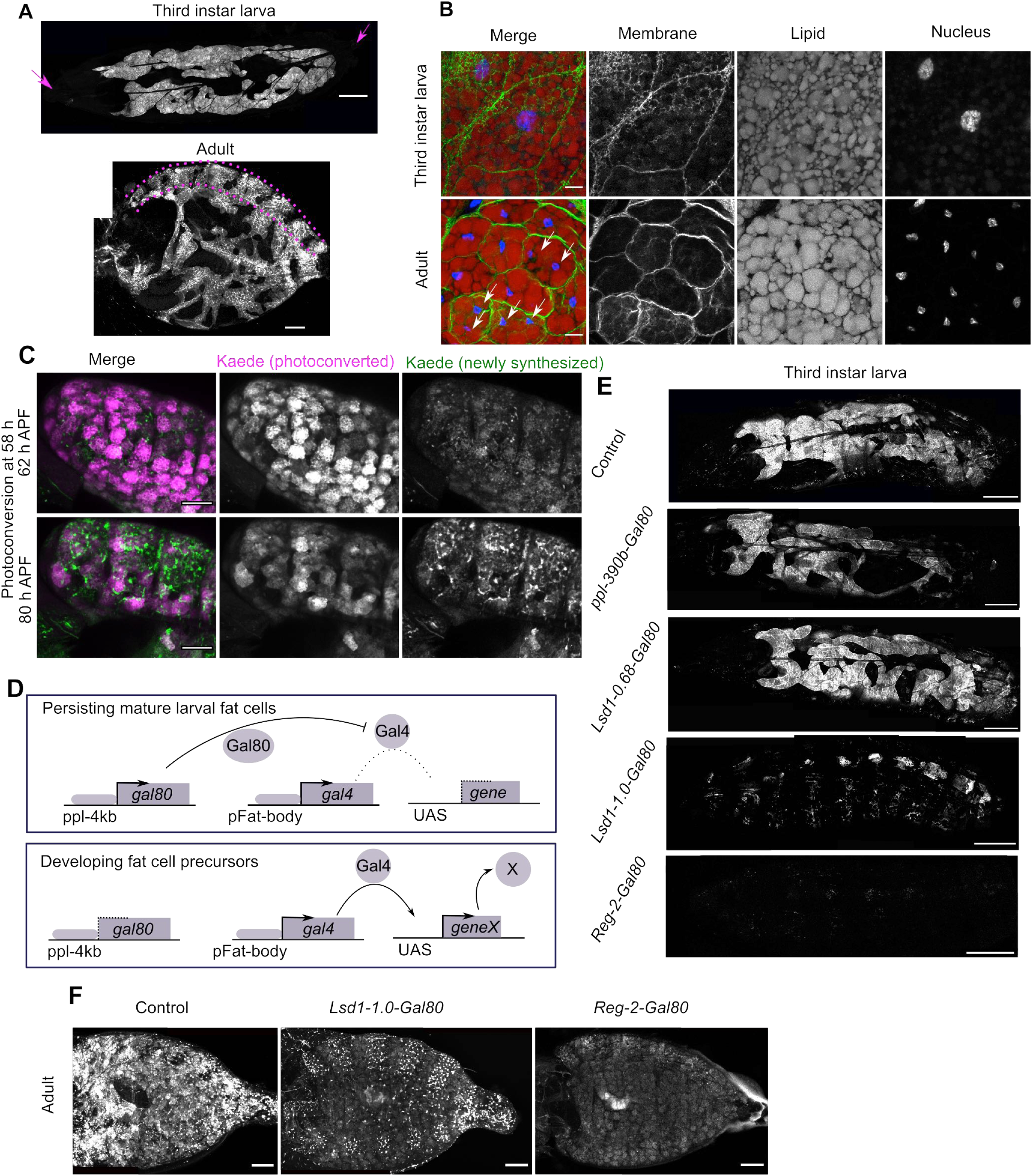
Several *Gal80* lines under the control of cis-elements of fat-body-related genes suppress Gal4-driven gene expression in the LFB and AFB. (A) Distribution of the LFB in a mature 3rd instar larva (top; dorsal view; arrows indicate the anterior and posterior ends) and the AFB in the abdomen at 7 days after adult eclosion (bottom; lateral view; the same image used in Fig.1A is shown for comparison). Fat bodies were visualized with membrane-bound GFP markers. (B) Numerous lipid droplets were deposited in LFB cells in a mature 3rd instar larva (top) and AFB cells at 5 days after adult eclosion (bottom). Note that LFB cells have a single large polyploid nucleus and that AFB cells often contained multiple small nuclei (white arrows). (C) Kaede photoconversion revealed the AFB in a pharate adult fly. For photoconversion, a fly expressing Kaede driven by *c833-Gal4* was illuminated with UV light at 58 h APF. Newly synthesized, unconverted green fluorescence signals were predominantly observed in adult fat cells and highlighted the AFB at 80 h APF. (D) Schematic illustrations showing how *ppl-4kb-Gal80* suppresses Gal4 function in mature LFB cells but not in developing adult fat precursor cells. Gal4 binds to the UAS- sequence and induces the expression of genes located downstream of the sequence, and Gal80 inhibits the Gal4-dependent gene expression. The genomic element utilized for the *ppl-Gal4*, a widely used fat-body Gal4, was inserted as the upstream cis-element for our *Gal80* construct. The *ppl* gene has been reported to express in later stages of LFB development during embryogenesis (Zinke et al., 1999). *ppl-4kb-Gal80* is designed to act as an off-switch for Gal4 under the control of fat-body-related cis-elements (pFat-body) in persistent, mature larval fat cells but not in developing adult fat cells. (E-F) Fat body-related *Gal80* genes (see Fig. S2A-D and Table 1) established in this study suppressed *Cg-Gal4*-driven GFP expression in fat cells*. Lsd-1.0kb-Gal80* and *Reg-2-Gal80* diminished GFP signals in the LFB of mature larvae (E). The suppression by these *Gal80* transgenes was also observed in the LFB and the AFB of pharate adult flies shortly before eclosion (F). Scale: 500 μm (A top, E), 200 μm (A bottom, F), 10 μm (B), 100 μm (C).

**Fig. S2.**
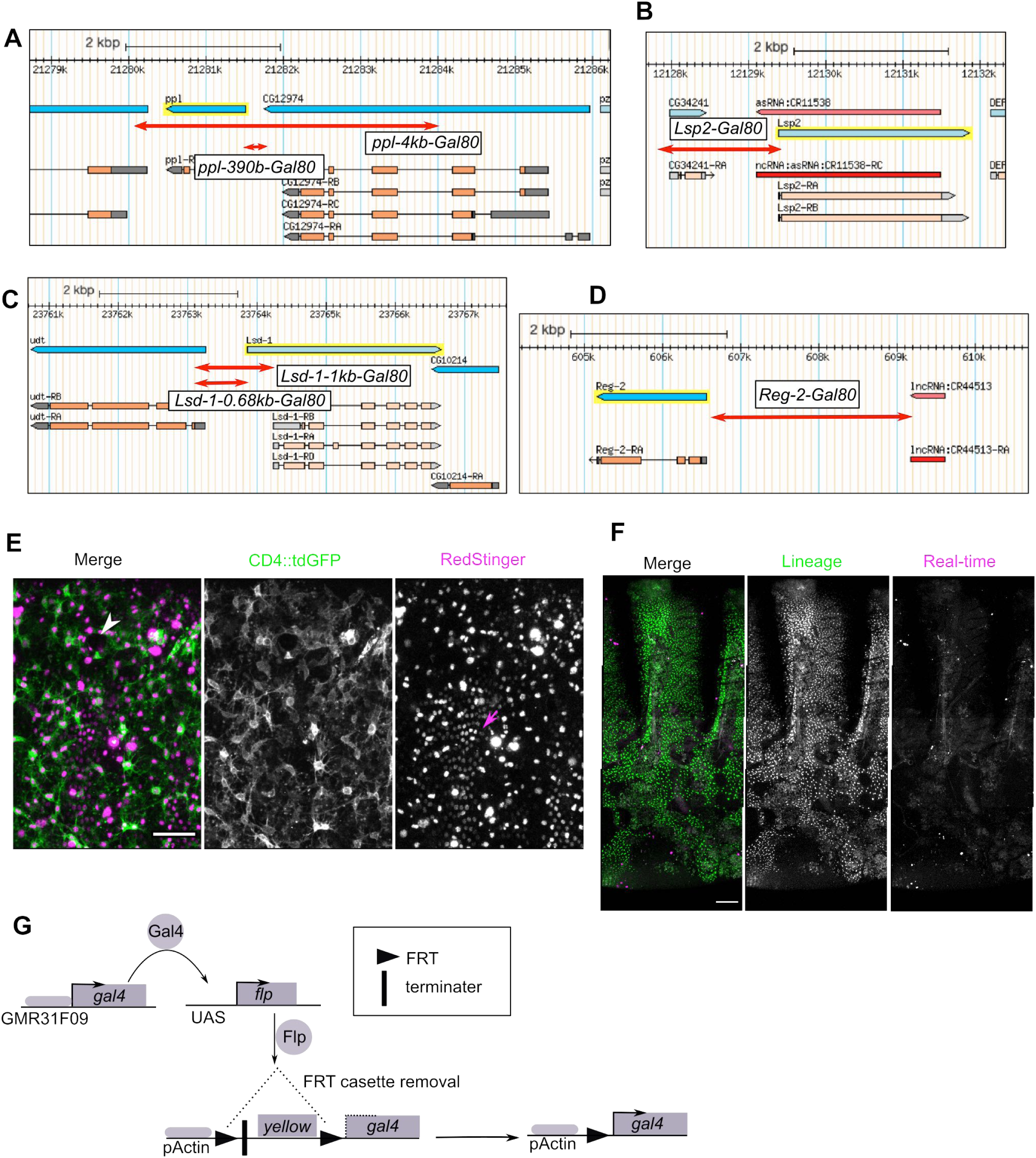
Genomic tools for Gal4-dependent gene expression in the AFB lineage. (A-D) Regions of cis-elements on the fly genome used for generating fat-body *Gal80* strains are indicated by red lines with double arrowheads on captured images from the GBrowse viewer at Flybase (Thurmond et al., 2018; screenshots were captured on March 10, 2018). (E) *c833-Gal4* plus *ppl-4kb-Gal80* induced marker expression in non-AFBp cells as well as AFBp cells. Lateral view images of the pleurite of a fly expressing CD4::tdGFP and RedStinger, nuclear-targeted DsRed, at 48 h APF. The white arrowhead indicates the abdominal spiracle. Red Stinger signals were not specific in AFBp cells. For instance, regularly arranged cells (magenta arrow), which might be a subset of adult epidermal cells, exhibited weak RedStinger signals. (F) Transient expression of *svp[31F09]-Gal4* in an adult fat cell lineage was revealed using G-TRACE in a newly eclosed adult female fly (Evans et al., 2009; see Materials and Methods). Stinger signals (Lineage) but not RedStinger signals (Real-time) were seen in the AFB in the presence of *svp[31F09]-Gal4*, suggesting a transient expression early in AFB development. (G) *Ay-Gal4*-based persistent labeling of AFBp cells after transient Gal4 expression. In the *Ay-Gal4* construct, Gal4 expression driven by the actin promoter (pActin) is interrupted by an upstream transcriptional terminator within an FRT cassette containing the *yellow* gene (Ito et al., 1997). Transient Gal4 expression driven by *svp[31F09]-Gal4* (*GMR31F09*) activates *UAS-FLP*, resulting in the reconstitution of Actin-Gal4 in the lineage by excising the FRT cassette. Scale: 50 μm (E) and 100 μm (F).

**Fig. S3.**
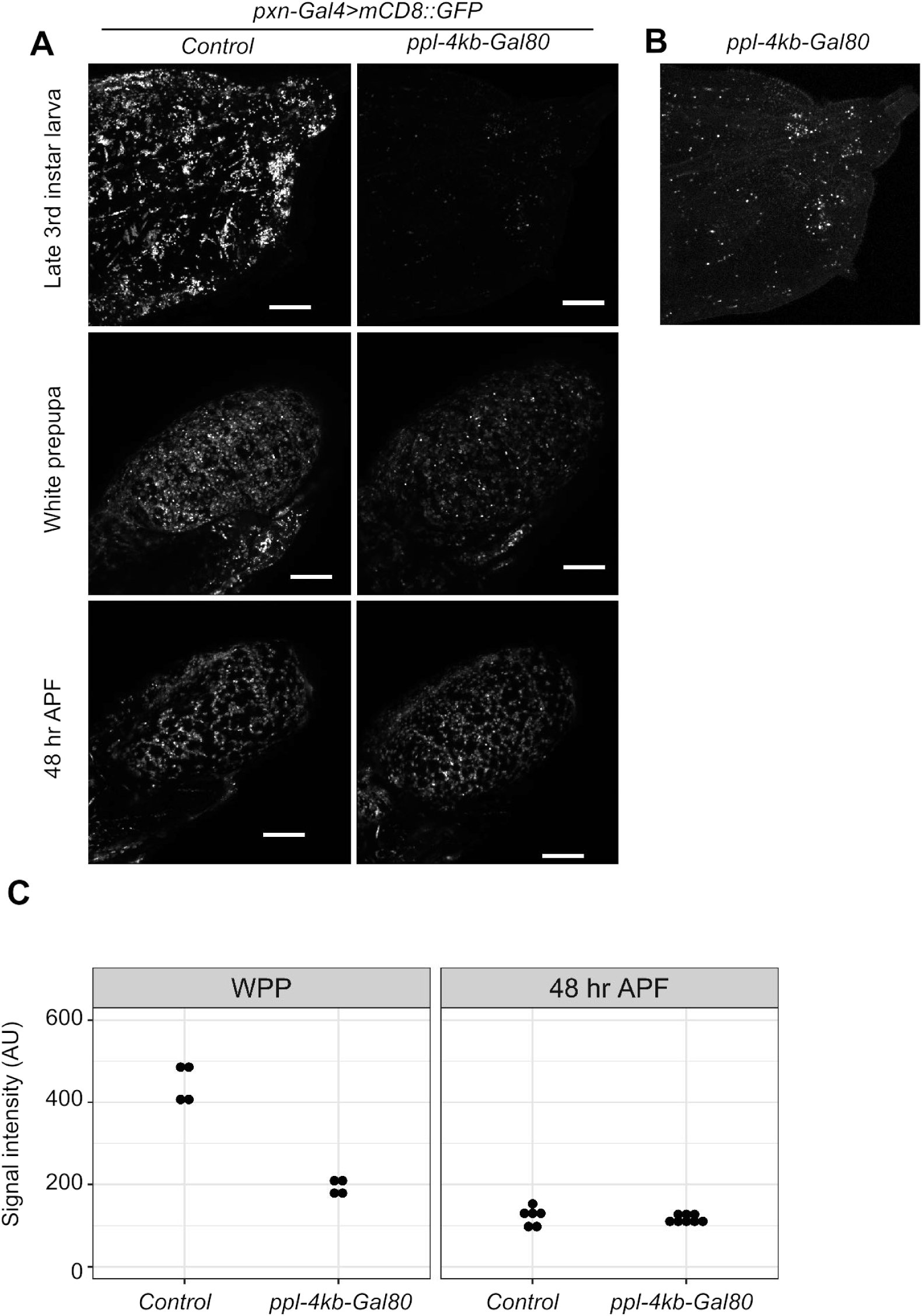
p*p*l*-4kb-Gal80* suppresses *pxn-Gal4*-driven marker expression in hemocytes. (A-C) *ppl-4kb-Gal80* lowered GFP signals in hemocytes (*pxn-Gal4>mCD8::3xEGFP*) at the mature 3rd instar stage and immediately after puparium formation (the white prepupa stage), but not at 48 h APF. (A-B) Representative fluorescent images of the posterior end of a mature larva (A, top) and the abdomen of a white prepupa (A, middle) and a pharate adult fly at 48 h APF (A, bottom). *ppl-4kb-Gal80* strongly suppressed GFP signals in the late 3rd instar stage (A, top). The brightness of the top-right image in (A) was adjusted to highlight weak GFP signals in hemocytes (B). (C) Quantification of average GFP signal intensities of the abdomen at the white prepupa stage (WPP) and at 48 h APF. Each dot indicates a signal intensity value calculated from a single fly. The total intensity value of each abdomen in z-projected images was subtracted by background signals and then divided by the pixel number of the abdomen. Scale: 200 μm.

**Fig. S4.**
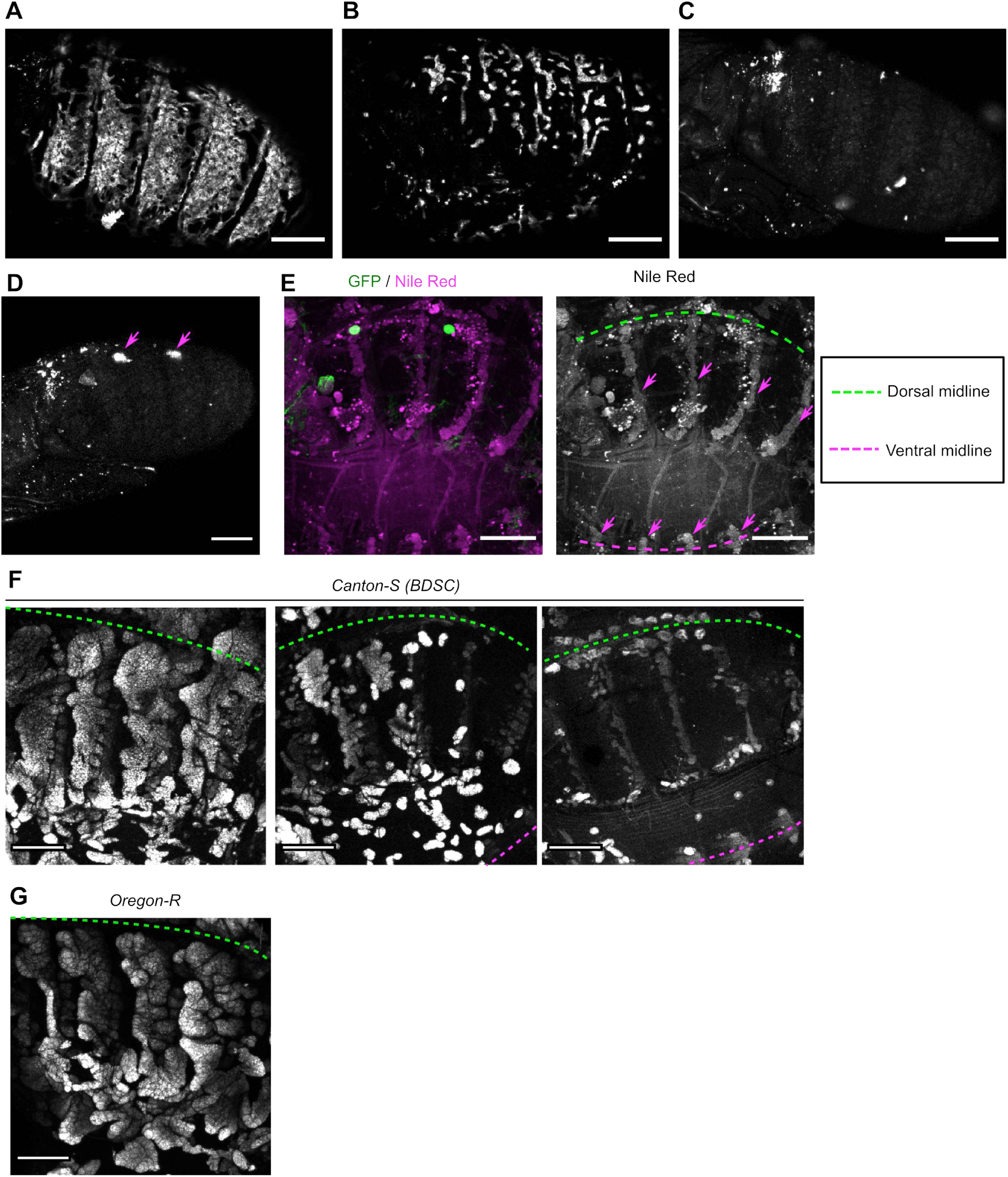
Wildtype flies occasionally display developmental defects in the AFB. (A-C) Pharate adult flies expressing CD4::tdGFP in the AFB (*svp[31F09]+Ay>CD4::tdGFP*) showed developmental defects to various degrees (70-80 h APF: A, fully developed; B, partially disrupted; C, almost completely lost; see also Table 2). (D-E) AFB defects in pharate adult flies with reduced AFB markers were confirmed with Nile Red staining (N = 11 out of 11 flies tested). Whole-mount live imaging revealed GFP signals in non-AFB cells, including nephrocytes (arrows in D). Nile Red staining of the fixed fillet preparation of the same fly showed the absence of AFB but highlighted oenocytes with weak signals (arrows in E). (F,G) *Canton-S*, but not *Oregon-R*, occasionally exhibited partially or almost complete loss of Nile Red signals, indicating developmental defects of the AFB (estimated defect probability: approximately 30%; see also Table 3). Scale: 200 μm.

**Fig. S5.**
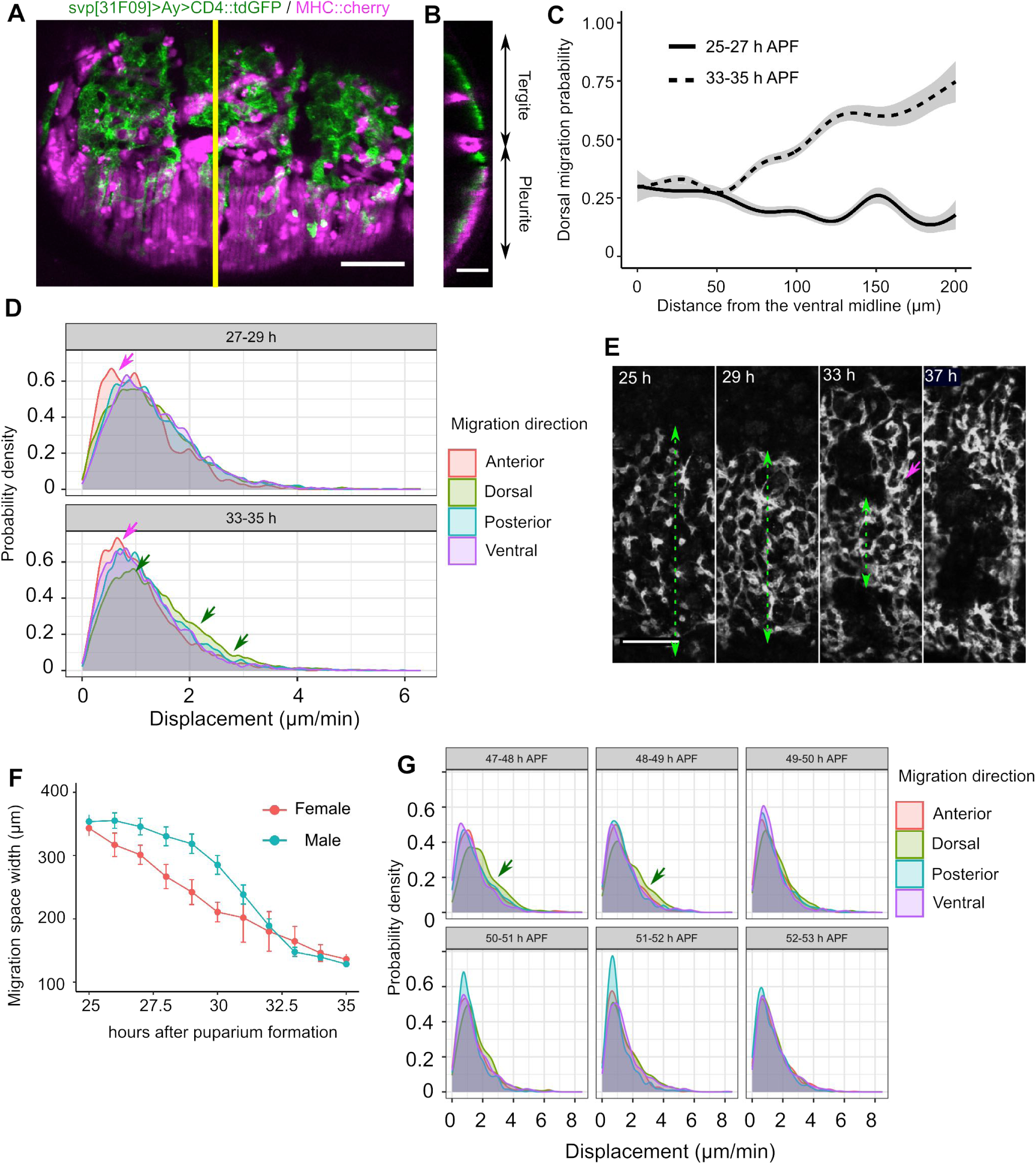
AFBp cells migrate in a temporally regulated and directionally oriented manner in the abdomen. (A, B) The AFB resides at superficial levels within the abdomen. (A) Lateral view image of the AFB and muscle (73 h APF) visualized with CD4::tdGFP and the endogenous MHC proteins tagged with mCherry (MHC::cherry), respectively. (B) The y-z orthogonal image along the yellow line in (A), with the deeper part (the body cavity) to the left. The adult fat cells were located below lateral muscles in the pleurite and above retractor muscles in the tergite. AFBp cells migrate at the same superficial levels in the abdomen and are readily observed by fluorescence microscopy without dissection. (C,D) Quantification of the dorsal migration probability and displacement values of ventral AFBp cells. Estimated probability that a given observed step (> 2 μm/2 min) is toward the dorsal direction (45°-135° in Fig. 2E) as a function of distance from the VML (C). Dorsal migration probabilities were estimated using generalized additive models (N = 5168 and 9411 steps from 4 flies for 27-29 h and 33-35 h, respectively). Shaded areas indicate 2 x standard error. Density plots show migration displacement values split by developmental periods and directional categories (D; N = 8958 and 17383 steps in total for 27-29 h and 33-35 h AFP, respectively). Anterior movements (magenta arrows) tended to be slightly smaller than movements toward other directions in both periods. At 33-35 h APF, the distribution of dorsal movements peaked at a slightly larger value and had a higher right-side tail (green arrows) than migrations toward other directions. (E,F) Representative image sequences (E) and quantification (F) of progressive decreases in the migration area width of ventral AFBp cells (dashed green lines with arrowheads). Arrow indicates a path for dorsally migrating AFBp cells. Error bars indicate standard deviation (N = 6 female and 8 male flies). (G) Displacement values during 47-53 h APF split by developmental periods and directional categories (G; N = approximately 2000 movements from 80 cells randomly selected from 4 flies for each time-window). Migrations toward the dorsal direction tended to be larger than migrations toward other directions during 47-49 h APF. Scale: 100 μm (A, E) and 50 μm (B).

**Fig. S6.**
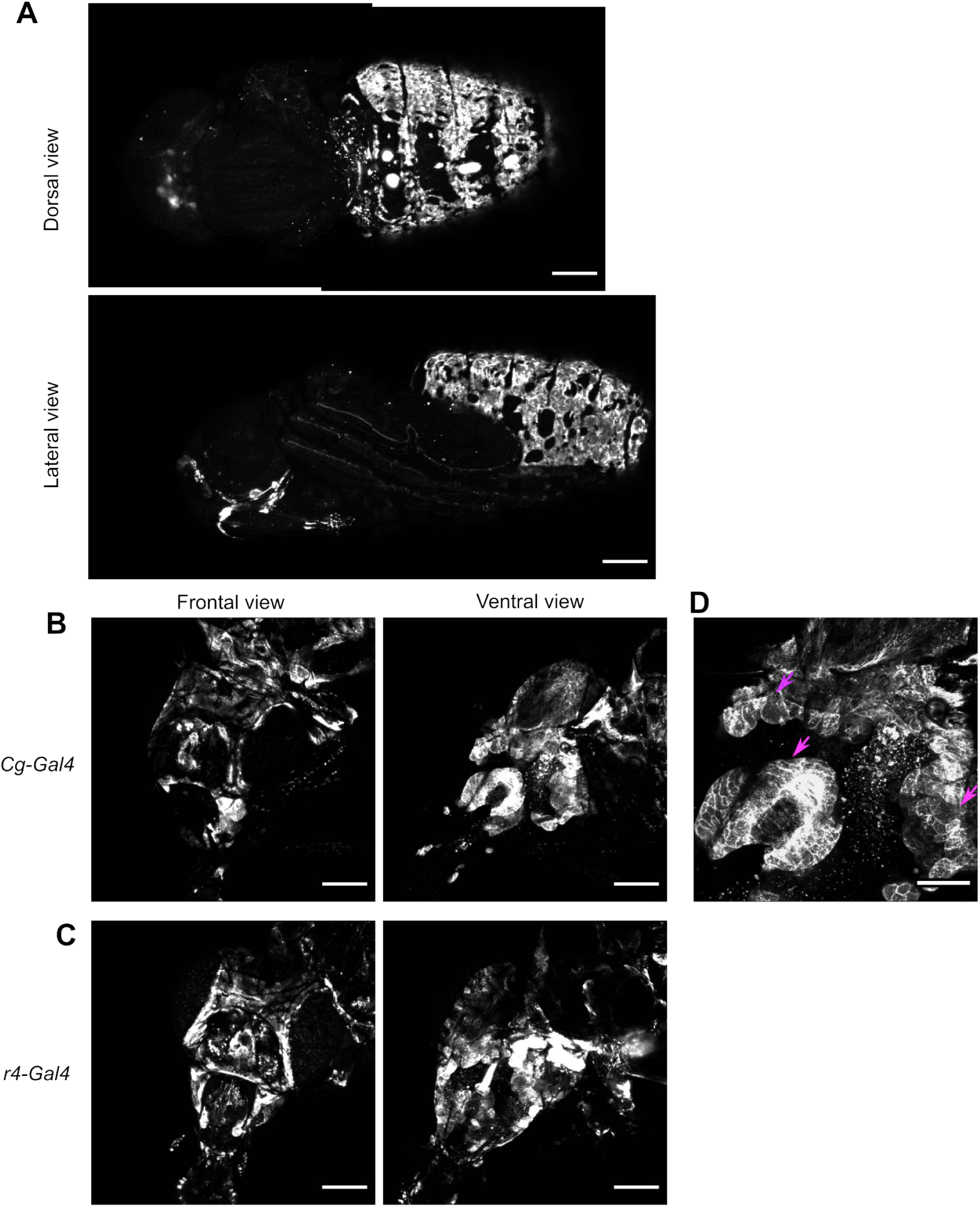
The AFB in the adult head. (A) Representative images of flies expressing CD4::tdGFP driven by *svp[31F09]-Gal4* plus *Ay-Gal4* (73 h APF) with strong GFP signals in the abdominal AFB but not in the head AFB. *svp[31F09]-Gal4* plus *Ay-Gal4* expression might be specific in abdominal populations (N = 7). (B-D) Fluorescent images of seven-day-old female fly heads expressing mCD8::GFP under the control of *Cg-Gal4* (B) and *r4-Gal4* (C), widely used fat body Gal4 strains. These *Gal4* drivers showed similar expression patterns in the head. Most GFP-positive cells showed a sheet-like arrangement (arrows in D, higher magnification of image in B). Both *Cg-Gal4* and *r4-Gal4* were strongly expressed in the abdominal AFB at the same age (data not shown). Scale: 200 μm (A-C), 100 μm (D)

**Fig. S7.**
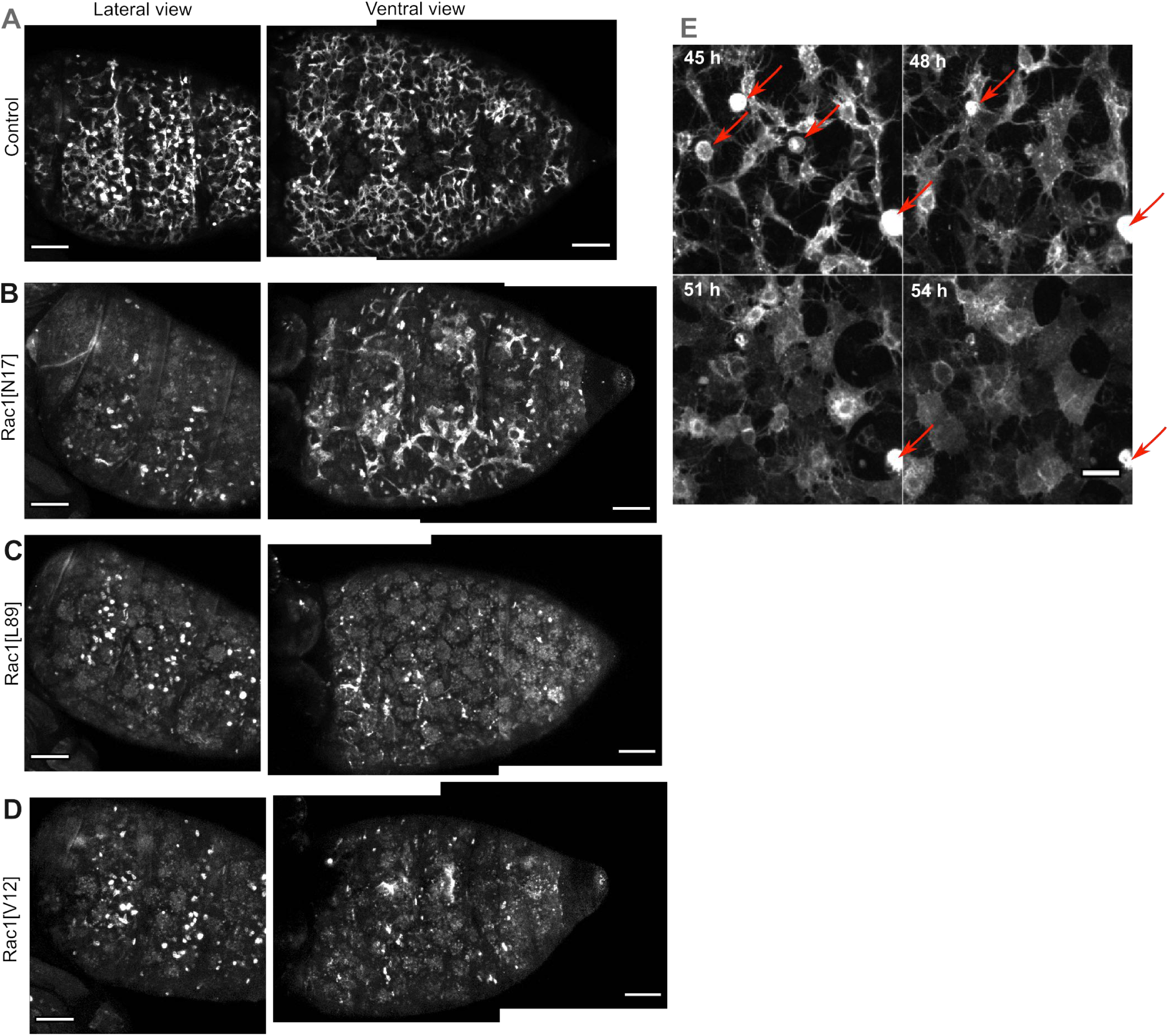
Ectopic expression of Rac1 mutants disturbs AFBp cell dynamics in the abdomen. (A-E) Representative images of the abdomen of flies expressing dominant-negative or constitutively active mutant forms of Rac1 (approximately 48 h APF at 29°C, see Table 11). *c833-Gal4* plus *ppl-4kb-Gal80* in conjunction with *tub-Gal80[ts]* was adopted for timed induction of Gal4-driven expression after puparium formation (McGuire et al., 2004). Rac1[N17] and Rac1[L89] are dominant-negative mutants, and Rac1[V12] is a constitutively active form (Luo et al., 1994). GFP positive cells in lateral views of Rac1[N17], Rac1[L89], and Rac[V12] are not likely to be AFBp cells, as judged by their shape and dynamics. These cells were also found in control backgrounds (E, arrows), exhibited a spherical shape, and migrated to the deeper levels of the body cavity over time. Scale: 100 μm (A-D), 10 μm (E).

**Fig. S8.**
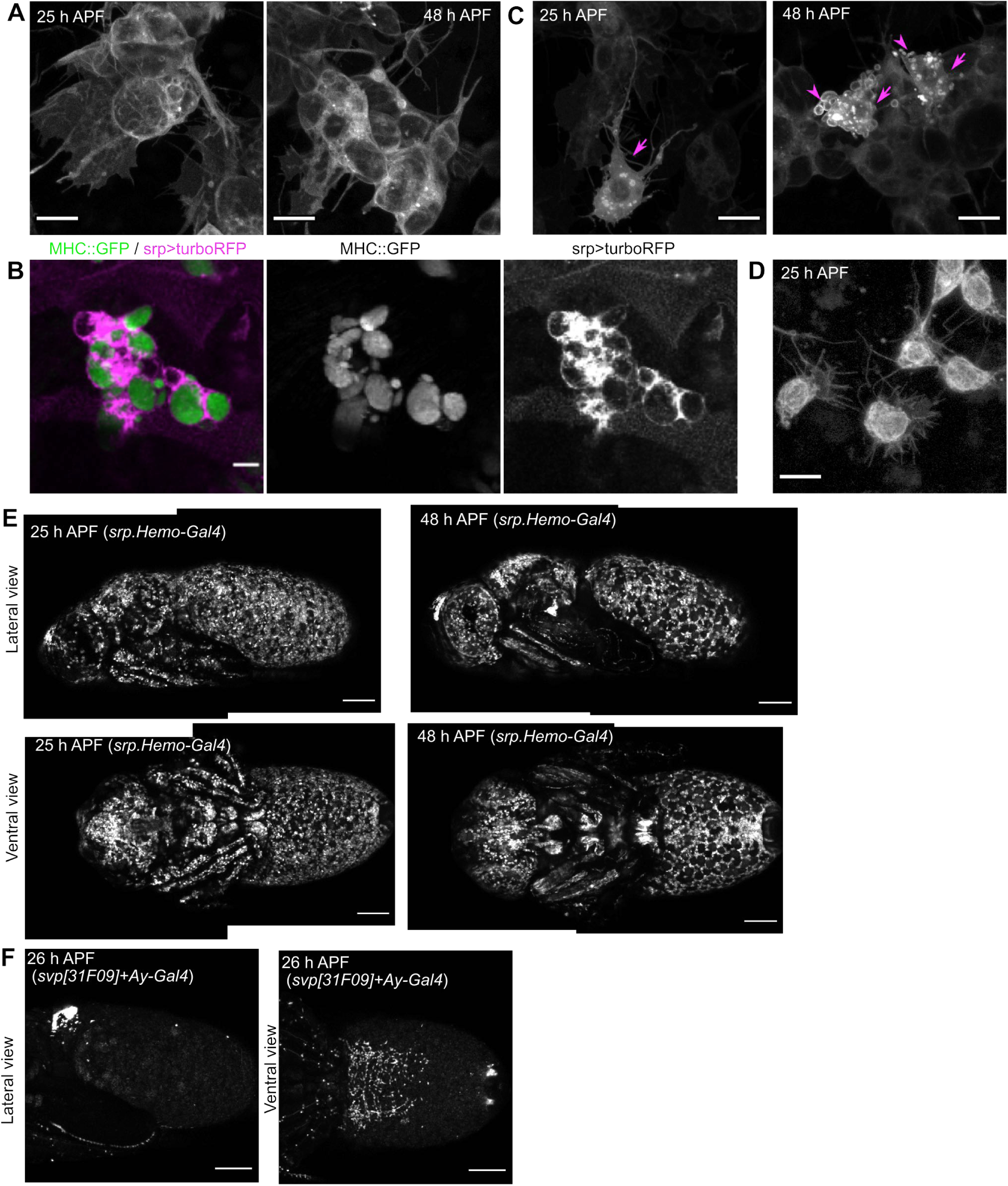
Morphologies and distributions of hemocytes during metamorphosis are distinct from those of AFBp cells. (A-D) Cell morphologies of hemocytes and AFBp cells. The vast majority of cells expressing *srp.Hemo-Gal4* were plasmatocytes characterized by prominent lamellipodia-like protrusions and intracellular vacuoles (A, see also Movie 21). These vacuoles were often colocalized with MHC::GFP signals, suggesting that they contained histolyzed larval tissues, including debris derived from larval muscle cells (B; Ghosh et al., 2020). Another class of *srp.Hemo-Gal*4-expressing cells consists of putative crystal cells (C; arrows). These cells lacked vacuole-like inclusions and often showed numerous bulges on their surface at 48 h APF (arrowheads). Both types of hemocytes were morphologically distinct from AFBp cells (D; membrane GFP signals nonlinearly adjusted to highlight fine protrusions). (E-F) Distributions of hemocytes and AFBp cells in the fly. Hemocytes were distributed across the whole body of flies at 25 and 48 h APF (E; *srp.Hemo-Gal4>mCD8::GFP*). AFBp cells were only seen in the anterior regions of the ventral side at 26 h APF in the abdomen (F; *svp[31F09]-Gal4* plus *Ay-Gal4>mCD8::GFP*). In the lateral view image (F), GFP signals at the dorsoanterior end of the abdomen would be debris of larval cells that had expressed *svp[31F09]-Gal4*. Scale: 10 μm (A-D), 200 μm (E, F).

**Fig. S9.**
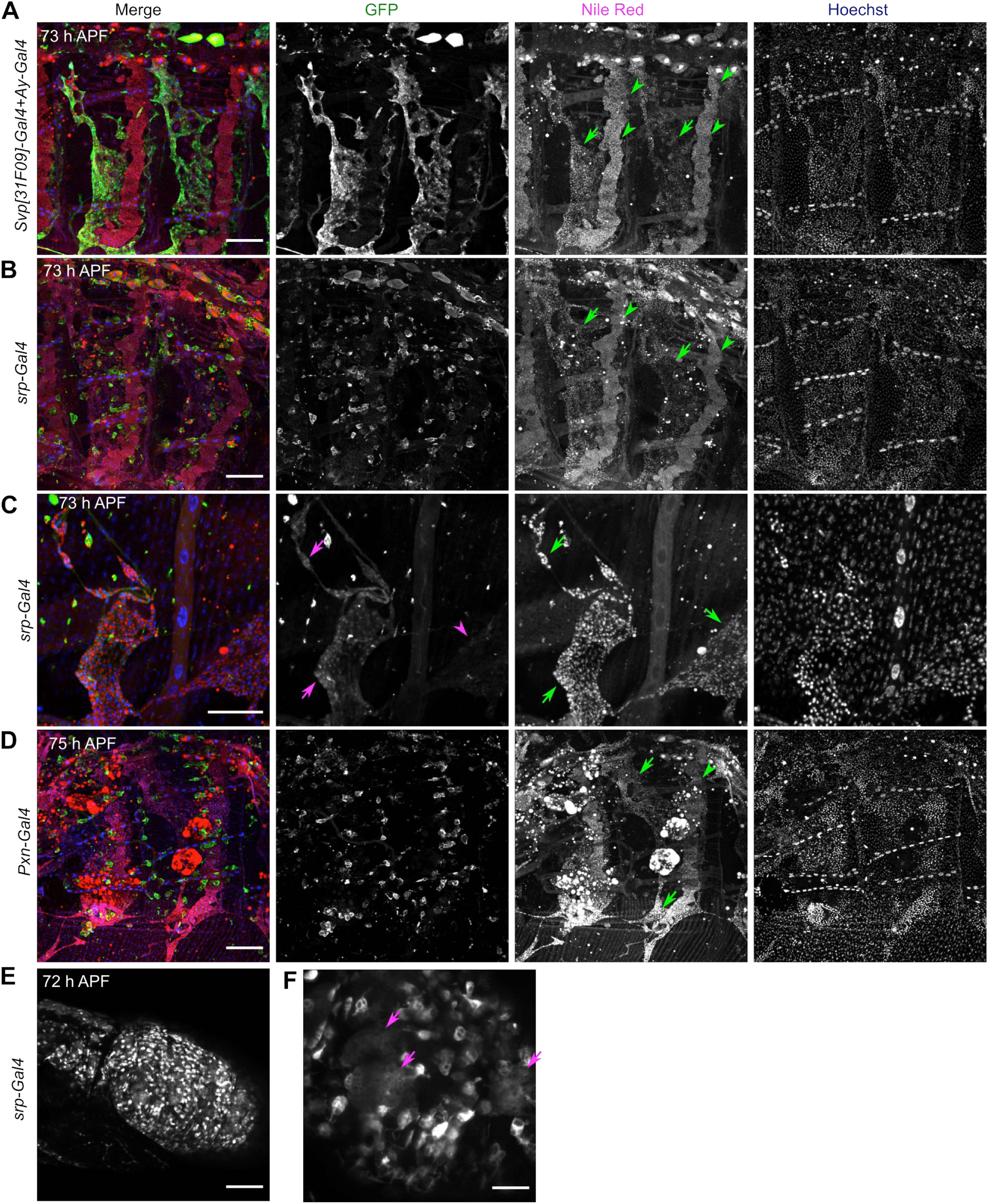
Cells derived from the hemocyte lineage may not directly contribute to the AFB. (A-D) Representative images of pharate adult flies expressing GFP markers in the AFB (A: *31F09-Gal4* plus *Ay-Gal4>CD4::tdGFP*) or hemocytes (B,C: *srp.Hemo-Gal4> mCD8::GFP*; D: *pxn-Gal4>CD4::tdGFP*). Pharate adult flies at indicated hours after puparium formation were dissected and stained with Nile Red plus Hoechst 33342 and immuno-stained using GFP antibody to augment GFP signals (A-D). The tergite is shown in (A), (B), and (D). A higher magnification image of part of the pleurite is shown in (C). Nile Red signals were mainly detected in oenocyte belts (green arrowheads) and the AFB (green arrows). GFP signals driven by *31F09-Gal4* plus *Ay-Gal4* were colocalized with Nile Red Signals (A). The AFB in flies expressing GFP induced by hemocyte Gal4s almost completely lacked GFP signals (B, D; N = 3 for *srp.Hemo-Gal4* and *pxn-Gal4*, respectively). In a single fly with *srp.Hemo-Gal4*-driven GFP expression, part of the AFB displayed weak GFP signals (magenta arrows in the GFP channel of C), but other AFB regions lacked GFP signals (magenta arrowhead in the GFP channel of C). (E-F) *srp.Hemo-Gal4* was expressed in the LFB during metamorphosis. Large, spherical larval fat cells were seen at 72 h APF (arrows in F). GFP signals were detected from 45 h APF onward in *srp.Hemo-Gal4>UAS-mCD8::GFP* flies. Scale: 100 μm (A, B, D), 50 μm (C, F), and 200 μm (E)

**Fig. S10.**
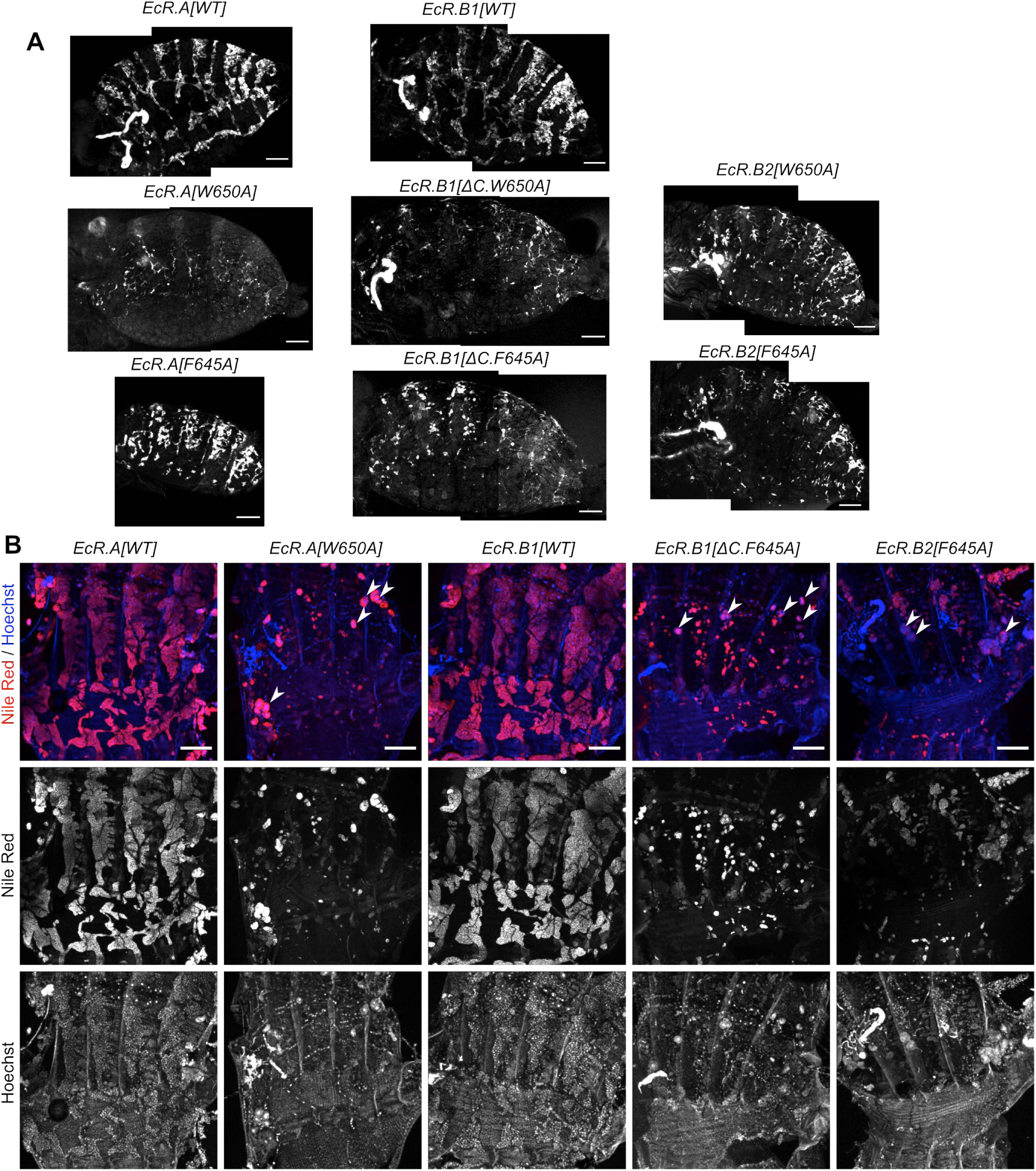
Attenuation of ecdysteroid-dependent gene activation disrupts AFB at late developmental stages. (A) Representative images of the abdominal AFB overexpressing wild-type ecdysone receptors and dominant-negative forms of ecdysone receptors. EcR.A[W650A], EcR.A[F645A], EcR.B1[ΔC.W650A], EcR.B1[ΔC.F645A], EcR.B2[W650A], and EcR.B2[F645A] are dominant-negative forms of EcR. Transgene activation was induced after puparium formation using *C833-Gal4* plus *ppl-4kb-Gal80* in conjunction with *tub-Gal80[ts]*, and flies were observed at 75 h APF, after incubation at 29°C. The fully developed AFB was seen in *EcR.A[WT]* and *EcR.B1[WT]*-expressing flies. Large GFP-positive tissues at anterior regions corresponded to be the adult salivary gland. (B) Representative images of the AFB overexpressing wild-type ecdysone receptors and dominant-negative forms of ecdysone receptors stained with Nile Red. Flies were aged at 29°C for 100 hours. White arrowheads indicate persistent larval fat cells, as judged by the presence of a large nucleus in the Hoechst channel. See also Table14 and Movie 22-23. Scale: 200 μm.

**Fig. S11.**
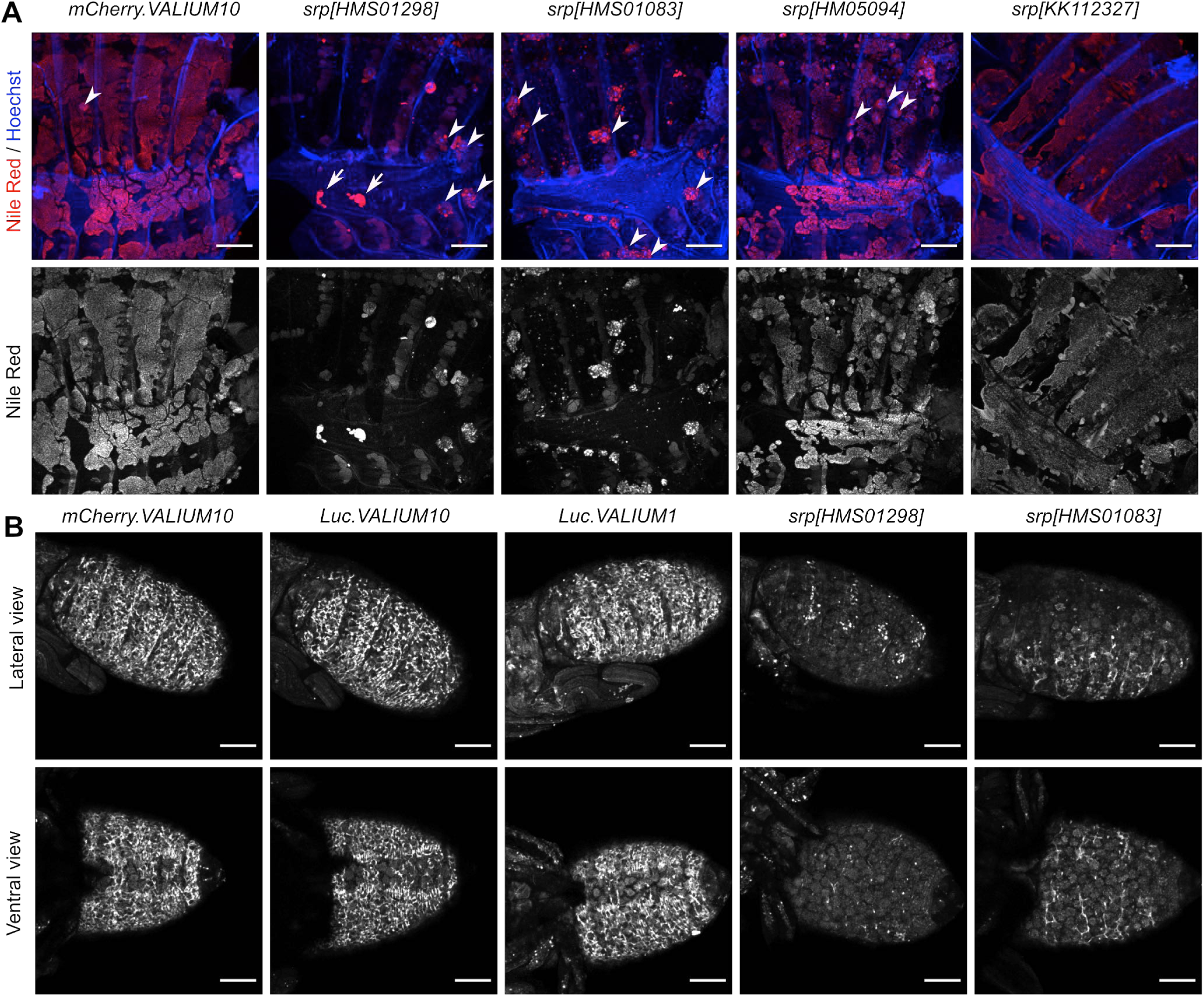
RNAi-mediated silencing of *srp* causes thread-like cellular assemblies. (A) Representative images of the AFB in flies ubiquitously expressing RNAi transgenes targeting *mCherry* (control) or *srp*. These flies were grown at 19°C until 0-6 h APF. The RNAi transgenes were expressed in the whole body by *tub-Gal4* in conjunction with *tub-Gal80[ts]* by the temperature induction from 0-6 h APF onward. Flies were dissected at approximately 100 h APF and stained with Nile Red, revealing that *srp* knockdown by *srp[HMS01298]* and *srp[HMS01083]* caused an almost complete loss of the AFB. White arrows indicate AFB cells, and white arrowheads indicate persistent larval fat cells, as judged by the presence of a large nucleus in the Hoechst channel. (B) Representative images of AFBp cells expressing RNAi transgenes targeting *mCherry* (control)*, Luc* (control), or *srp.* Silencing was induced after puparium formation using *C833-Gal4* plus *ppl-4kb-Gal80* in conjunction with *tub-Gal80[ts]*, and flies were observed at 50 h APF, after incubation at 29°C. See also Table 15 and Movie 24-27. Scale: 200 μm (A and B).

## Movie legends

Movie 1

AFBp cells were distributed in a quasi-two-dimension manner in the abdomen.

Representative 3D rotate image of AFBp cells expressing CD4::tdGFP (cell membrane, green) and RedStinger (nucleus, magenta) driven by *31F09-Gal4* plus *Ay-Gal4* at the pleurite (47 h APF). Numbers in the movie refer to the scale in micros. The z-axis indicates depth in z-series image sequences, with smaller numbers to the body cavity. Anterior is to the left, and dorsal is up in the first frame of the movie.

Movie 2

Migrating AFBp cells extend lamellipodia-like and filopodia-like protrusions.

Representative time-lapse images of migrating AFBp cells expressing CD4::tdGFP and RedStinger driven by *31F09-Gal4* plus *Ay-Gal4* at 47 h APF. The cell membrane and nucleus were labeled with CD4td::GFP (green, middle) and RedStinger (magenta, right), respectively. Anterior is to the left, and dorsal is up.

Movie 3

AFBp cells emerge from the ventroanterior end of the abdomen and then migrate posteriorly.

Representative ventral view movie of AFBp cells expressing CD4::tdGFP driven by *31F09-Gal4* plus *Ay-Gal4* (13-27 h APF). At first, some axonal projections extending from the central nervous system were seen and then underwent metamorphic degeneration. AFBp cells appeared at 15 h APF and then migrated posteriorly while they frequently divided. Anterior is to the left.

Movie 4

AFBp cells cover the whole ventral abdomen by 40 h APF.

Representative ventral view movie of AFBp cells expressing CD4::tdGFP driven by *31F09-Gal4* plus *Ay-Gal4*. At first, AFBp cells accumulated around the ventral midline and switched their migration direction to cover the entire region of the ventral abdomen. The marker also highlighted some cells around the genitalia, and the male-specific genitalia rotation was observed from 28 to 33 h APF (Inatomi et al., 2019). Anterior is to the left.

Movie 5

A subset of ventral AFBp cell populations start migrating dorsally at approximately 30 h APF.

Representative ventral view movie of AFBp cells expressing Stinger driven by *31F09-Gal4* plus *Ay-Gal4* in the abdomen (A3-4 segments). Cells with slightly larger nuclei are likely to be those of hemocytes. Anterior is to the left.

Movie 6

The migration area for ventral AFBp cells progressively narrows.

Representative ventral view movie of AFBp cells expressing CD4::tdGFP driven by *31F09-Gal4* plus *Ay-Gal4* in the abdomen. While AFBp cells migrated posteriorly, they progressively concentrated along the ventral midline, with a subset of cells starting dorsal migration toward the pleurite. Anterior is to the left.

Movie 7

Ventral AFBp cells start dorsal migration from 30 h APF onward.

Representative lateral view movie of abdominal AFBp cells expressing CD4::tdGFP driven by *31F09-Gal4* plus *Ay-Gal4*. Until 30 h APF, AFBp cells were only seen at the ventral side of the abdomen and migrated in the posterior direction. After 30 h APF, a subset of these cells started moving dorsally and covered almost the entire areas of the pleurite and tergite. Anterior is to the left, and dorsal is up.

Movie 8

AFBp cells from the ventral side of the abdomen reached the dorsal midline.

Representative dorsal view movie of the anterior part of the abdomen of a fly expressing CD4::tdGFP driven by *c833-Gal4* plus *ppl-4kb-Gal80* in AFBp cells. AFBp cells entered the tergite by 43 h APF and reached the dorsal midline by 50 h APF. Anterior is to the upper left. Scale: 100 μm.

Movie 9

AFBp cells at the pleurite reverse their migration direction during 47-53 h APF.

Representative lateral view movie of abdominal AFBp cells expressing a nuclear GFP marker driven by *31F09-Gal4* plus *Ay-Gal4* (A2-A3 segments). The arrow and arrowheads in the first frame indicate the abdominal spiracle and immotile cells around the spiracle, respectively. By 49 h APF, AFBp cells at regions above the spiracle (dorsal to the spiracle) outnumbered those at regions below the spiracle (ventral to the spiracle). Their migration direction gradually reversed toward the ventral direction, resulting in a more uniform distribution of AFBp cells in the lateral part of the abdomen by 53 h APF. Anterior is to the left, and dorsal is up.

Movie 10

Hemocytes and larval fat cells at the pleurite are almost immotile during 47-53 h APF.

Representative lateral view movie of hemocytes and larval fat body cells expressing a nuclear GFP marker (*srp.Hemo>Stinger*) during 47-53 h APF (A2-A3 segments). Small and large Stinger signals represented nuclei of hemocytes and larval fat cells, respectively. Their movements were distinct from those of AFBp cells in the same region during the same developmental period (Movie 9). Anterior is to the left, and dorsal is up.

Movie 11

AFBp cells dispersed throughout almost the entire abdomen and then assemble into the adult fat body.

Representative lateral view movie of the abdomen of a fly expressing CD4::tdGFP driven by *31F09-Gal4* plus *Ay-Gal4* in AFBp cells. AFBp cells ceased their migration by 60 h APF and then formed the sheet-like adult fat body by 65 h. Anterior is to the left, and dorsal is up.

Movie 12

AFBp cells form a sheet-like tissue by establishing lateral contacts with neighboring cells.

Representative high-magnification movie of AFB formation in the tergite by AFBp cells expressing CD4::tdGFP driven by *31F09-Gal4* plus *Ay-Gal4* during 56-72 h APF. At first, AFBp cells were highly polarized and extended lamellipodia-like and filopodia-like protrusions. By 60 h APF, their cell bodies became flattened while their fine protrusions partly remained. Flattened cells made lateral contacts with neighboring cells and merged into a sheet-like tissue, with their cell-cell boundaries becoming obscure. Anterior is to the left, and dorsal is up.

Movie 13

AFBp cells in the head migrated anteriorly.

Representative frontal view movie of AFBp cells in the head during 25-30 h APF, from a fly aged at 29°C from puparium formation onward. CD4::tdGFP expression were induced after puparium formation by using *c833-Gal4*, *ppl-4kb-Gal80,* and *tub-Gal80[ts]*. *tub-Gal80[ts]* was employed to suppress GFP signals in degrading larval salivary gland cells. AFBp cells predominantly migrated along the anteroposterior axis at the frons. Their migration steps were weakly biased toward the anterior direction, as quantified in Fig. 5 C. Anterior is to the left, with the right compound eye in the upper and the proboscis in the lower portion of the frame.

Movie 14

AFBp cells migrated anteriorly at the vertex of the head.

Representative dorsal view image sequences of AFBp cells in the head during 38-47 h APF, from a fly aged at 29°C. CD4::tdGFP in AFBp cells was induced as in Movie 13. AFBp cells emerged from the posterior end of the head and then migrated anteriorly. In the thorax, AFBp cells were rarely seen. Anterior is to the left.

Movie 15

Expression of a dominant negative Rac1 (Rac1[N17]) interrupts AFBp cell migration in the abdomen.

Representative lateral view movie of abdominal AFBp cells in the absence or presence of Rac1[N17] expression (the left and right movies, respectively). Rac1[N17] and CD4::tdGFP in AFBp cells were driven by *c833-Gal4*, *ppl-4kb-Gal80,* and *tub-Gal80[ts]*. Flies were aged at 29°C from puparium formation onward.

Movie 16

Expression of a constitutively active Rac1 (Rac1[V12]) mutant causes AFBp cell clumps at the anterior end of the abdomen.

Representative ventral view movie of abdominal AFBp cells in the absence (left) or presence (right) of Rac1[V12] expression. Rac1[V12] and CD4::tdGFP in AFBp cells were driven by *c833-Gal4*, *ppl-4kb-Gal80,* and *tub-Gal80[ts]*. Flies were aged at 29°C from puparium formation onward. Rac1[V12] expression resulted in immotile cellular aggregates, which were likely to be eventually moved into the body cavity. Note that GFP-positive cells were only seen in the anterior part of the abdomen. Anterior is to the left.

Movie 17

AFBp cells undergo cell division during their migration period.

Representative movie of a migratory AFBp cell undergoing cell division at 47 h APF. CD4::tdGFP (green, middle) and RedStinger (magenta, right) were driven by *31F09-Gal4* plus *Ay-Gal4*. A highly polarized, migrating cell (indicated by the white arrow in the merge channel) rounded up, with its RedStinger signals becoming diffuse during 18-22 min. The resulting sister cells became polarized by 50 min and subsequently moved into different directions.

Movie 18

Adult fat body cells continue to divide even after the formation of the sheet-like architecture.

Representative movie of successive cell division events during 61-75 h APF. The AFB at the tergite was visualized by expressing CD4::tdGFP (green, middle) and RedStinger (magenta, right) under the control of *31F09-Gal4* plus *Ay-Gal4*. Cell membranes visualized with CD4::tdGFP highlighted part of the AFB with their sheet-like arrangement. RedStinger signals revealed frequent cell division events in adult fat body cells. Note that the total area size of CD4::tdGFP signals was largely unchanged during observations (14 hours), resulting in smaller adult fat body cells in the last frame.

Movie 19

Neighboring adult fat body cells are often fused to generate multinucleated cells.

Representative movie of cell fusion events at the tergite during 71-79 h APF. CD4::tdGFP in adult fat body cells was driven by *31F09-Gal4* plus *Ay-Gal4*. Arrows indicate cell-cell boundaries that diminished in the subsequent frames. A large migrating cell is likely to be a hemocyte. By 70 h APF, the fly started a muscle contraction, resulting in significant shifts in the x-y coordinates over time. The shifts in time-series images were automatically corrected by the Linear Stack Alignment with SIFT function in the Fiji software (Lowe, 2004). Anterior is to the left, and dorsal is up.

Movie 20

Pupal hemocytes display a distinct distribution and migration patterns compared with AFBp cells.

Representative ventral view movie of abdominal hemocytes expressing CD4::tdGFP (*srp.Hemo+ppl-4kb-Gal80>CD4::tdGFP*) during 26-49 h APF. Hemocytes were distributed across the abdomen, whereas AFBp cells were only seen at the ventral side of the abdomen at 26 h APF (Fig. S8F). Anterior is to the left, and dorsal is up.

Movie 21

The motility of hemocytes decreases by 50 h APF.

Representative ventral view movie of abdominal hemocytes expressing mCD8::GFP (*srp.Hemo>mCD8::GFP*) during 31-50 h APF. Most of the GFP-positive cells were plasmatocytes, which contain numerous vacuole-like inclusions. Smaller cells with stronger GFP signals were likely to be crystal cells. Both types of hemocytes were highly motile until 45 h APF. Hemocytes appeared to wrap around large spherical cells (i.e., larval fat cells) from 40 h APF onward.

Movie 22. *EcR.A1[W650A]* expression disrupts AFB development.

Representative movie of abdominal adult fat cells at the tergite in the absence (left) or presence (right) of EcR.A1[W650A] (a dominant negative mutant of EcR.A1). EcR.A1[W650A] and CD4::tdGFP were induced after puparium formation by using *c833-Gal4*, *ppl-4kb-Gal80,* and *tub-Gal80[ts].* AFBp cells expressing EcR.A1[W650A] showed minor defects in their migration period. They ceased migration by 55 h APF (at 29°C) and, then, displayed profound atrophic changes.

Movie 23. *EcR.B1[ΔC655.W650A]* expression disrupts AFB development.

Representative movie of abdominal adult fat cells at the tergite in the absence (left) or presence (right) of EcR.B1[ΔC655.W650A] (a dominant negative mutant of EcR.B1). EcR.B1[ΔC655.W650A] and CD4::tdGFP were induced as in Movie 22. Atrophic phenotypes after migration, similar to AFBp cells expressing EcR.A1[W650A], were seen (see Movie 22).

Movie 24. *srp* silencing by *srp[HMS01298]* in AFBp cells causes thread-like cellular assemblies.

Representative lateral view movie of AFBp cells in the absence (left) or presence (right) of *srp[HMS01298]*. *srp[HMS01298]* and *CD4::tdGFP* were under the control of *c833-Gal4*, *ppl-4kb-Gal80,* and *tub-Gal80[ts]*. Since *srp[HMS01298]* expression immediately after puparium formation resulted in the almost complete loss of AFBp cells (Fig. S11 B), knockdown efficiency early in metamorphic development was limited by placing flies at 19°C until 24 h APF, as in this experiment. Time-lapse imaging was started 8 hours after upshifting to a restrictive temperature. AFBp cells expressing s*rp[HMS01298]* appeared to adhere to each other after the start of dorsal migration and eventually formed thread-like cellular assemblies.

Movie 25. *srp* silencing by *srp[HMS01298]* in AFBp cells causes thread-like cellular assemblies (high-magnification).

Representative high-magnification lateral view movie of abdominal AFBp cells in the absence (left) or presence (right) of *srp[HMS01298]*. *srp[HMS01298]* and *CD4::tdGFP* were under the control of *c833-Gal4*, *ppl-4kb-Gal80,* and *tub-Gal80[ts]*. *srp[HMS01298]* expression was induced 24 h APF (19°C) as in Movie 24. *srp[HMS01298]*-expressing AFBp cells failed to be uniformly distributed and assembled into strikingly elongated cellular structures at 50 h APF.

Movie 26. *srp* silencing using *srp[HMS0183]* in AFBp cells caused web-like cellular assemblies.

Representative lateral view movie of abdominal AFBp cells in the absence (left) or presence (right) of *srp[HMS01083]* expression. *srp[HMS01083]* and *CD4::tdGFP* were driven by *c833-Gal4*, *ppl-4kb-Gal80,* and *tub-Gal80[ts].* Flies were aged at a restrictive temperature after puparium formation. AFBp cells expressing s*rp[HMS01083]* formed web-like networks by 40 h APF.

Movie 27. *srp[HMS01083]* expression in AFBp cells causes web-like cellular assemblies (high-magnification).

Representative high-maginification lateral view movie of abdominal AFBp cells in the absence (left) or presence (right) of *srp[HMS01083]* expression. *srp[HMS01083]* expression was induced 24 h APF, as in Movie 26.

Movie 28. Histoblasts cover the ventral side of the abdomen by 35 h APF.

Representative ventral view movie of abdominal histoblasts expressing mCD8::GFP (*esg-Gal4>Ay>mCD8::GFP*) during 26-38 h APF. Histoblasts expanded from the ventral nest to the ventral midline and covered the ventral side of the abdomen by 35 h APF (cf. Roseland and Schneiderman. 1979; Ninov et al., 2007). Oenocytes started showing strong GFP signals after 30 h APF.

Movie 29. Adult oenocytes establish their tissue arrangement by 35 h APF.

Representative lateral view movie of abdominal adult oenocytes expressing mCD8::GFP in a fly (*vvl[17C04]>mCD8::GFP*; see Materials and Methods for oenocyte Gal4 strains) during 20-44 h APF. Adult oenocytes started to express mCD8::GFP at approximately 25 h APF. Segmentally repeated oenocyte belts in the tergite were morphologically established by 35 h APF.

Movie 30. Simultaneous observation of AFBp cells and developing adult oenocytes.

Representative lateral view movie of abdominal AFBp cells and adult oenocytes, which were labeled with membrane-targeted GFP driven by *c833-Gal4*, *esg-Gal4*, *tub-Gal80[ts]* and *ppl-4kb-Gal80*. A fly was collected as a white prepupa (1 h APF) and grown at 19°C to suppress marker expression in undesired tissues (i.e., histoblasts and larval epidermis). Marker expression was induced at 25 h APF by upshifting to 29°C. Segmentally repeated oenocyte belts in the tergite were established prior to the start of the dorsal migration of ventral AFBp cells.

## Table title

Table 1 Summary of fat-body-related Gal80 constructs.

Table 2 The rate of disrupted AFB phenotypes in flies expressing fluorescent markers in the AFB.

Table 3 The rate of disrupted AFB phenotypes in wildtype strains stained with Nile Red.

Table 4 The rate of disrupted AFB phenotypes in DGRP strains.

Table 5 Quantification of ventral abdominal AFBp migration direction during 27-29 and 33-35 h APF.

Table 6 Quantification of ventral abdominal AFBp migration velocity during 27-29 and 33-35 h APF.

Table 7 Quantification of abdominal AFBp migration direction in the pleurite during 47-53 h APF.

Table 8 Quantification of abdominal AFBp velocity direction in the pleurite during 47-53 h APF.

Table 9 Quantification of head AFBp migration direction during 25-33 h APF (29°C).

Table 10 Quantification of head AFBp migration velocity during 25-33 h APF (29°C).

Table 11 The rate of disrupted AFB phenotypes in flies expressing mutant forms of Rac1.

Table 12 Quantification of the doubling time of AFBp cells in the ventral abdomen.

Table 13 Quantification of the doubling time of AFBp cells in the tergite.

Table 14 The rate of disrupted AFB phenotypes in flies expressing dominant-negative forms of EcR.

Table 15 The rate of disrupted AFB phenotypes in flies expressing RNAi constructs targeting Srp.

Table 16 Summary of fly strains and reagents used in this study.

Table 17 Summary of exact genotypes and sex of flies.

## Notes

### Competing Interest Statement

The authors have declared no competing interest.

